# Auditory Cortical Gradients Integrate Bottom-Up and Top-Down Structure During Natural Sound Categorisation

**DOI:** 10.1101/2025.11.18.689038

**Authors:** David Haydock, Robert Leech, Magdalena Kachlicka, Frederic Dick

**Affiliations:** Department of Experimental Psychology, University College London (UCL), UK; Institute of Psychiatry, Psychology & Neuroscience, King’s College London, UK; School of Psychological Sciences, Birkbeck, University of London, UK

**Keywords:** Auditory Cortex, fMRI, Functional Gradients, Representational Geometry, Spectral-Semantic Integration

## Abstract

Understanding how the brain organises natural categories is a central challenge in neuroscience. While prior work has shown that categories can be decoded from distributed activity patterns in auditory cortex, it remains unclear how these categories are globally arranged relative to one another, and how low-level acoustic and higher-level semantic structure jointly shape this organisation. Here, we addressed these questions by deriving low-dimensional functional gradients from high-depth functional magnetic resonance imaging (fMRI) data (three participants, ∼4.7 hours each) acquired during a category-specific one-back task. These gradients captured the principal axes of population activity in auditory cortex. Gradient-based models of the auditory cortex explained category structure more accurately than region-of-interest or whole-brain approaches, revealing that category information is distributed across multiple continuous axes rather than aligned with any single organisational dimension. Projecting acoustic (gammatone filter-bank) and behavioural similarity spaces directly into a shared framework with the fMRI functional axes showed that both contribute to the brain’s category geometry, with acoustic structure exerting a somewhat stronger influence. However, representational relationships varied across category pairs: some reflected primarily acoustic similarity, others semantic distinctions, and many a combination of both. This pairwise heterogeneity shows how auditory cortex may integrate multiple representational dimensions that define higher-level categories.

**Significance Statement:** Categorising natural sounds requires the brain to transform diverse acoustic signals into meaningful concepts. How this is achieved remains unclear: prior work has shown distributed activation patterns in auditory cortex, but not how category relationships are organised within its functional architecture. We show that natural sound categories are embedded across continuous cortical gradients that integrate bottom-up acoustic structure with top-down semantic information. Different categories emerge as flexible combinations of these spectral and behavioural dimensions. These findings reveal that natural sound categorisation arises from the geometry of cortical organisation, moving beyond localist accounts and offering a new framework for understanding how perception and cognition are linked in the human auditory system.

## 1 Introduction

Humans are remarkably adept at categorising the world around them. Categories—such as animals, tools, or emotions—are psychological groupings of stimuli that share common features. Even the concept of *sound* is a category, encompassing auditory events that share the property of being heard. How the brain constructs and represents such categories remains an open question despite extensive research (1), particularly for naturalistic stimuli like environmental sounds, where categories may be based on source identity (e.g., dog vs. bird), acoustic similarity (e.g., hum vs. drone), or personal associations (2).

In the auditory system, category structure appears to emerge as distributed population patterns rather than discrete “category units.” Early fMRI work showed that human superior temporal cortex carries reliable patterns that distinguish multiple natural sound categories (3). Studies since have shown that these neural patterns reflect not only acoustic similarity but also higher-level semantic or behavioural relationships between categories (4–8). However, most prior approaches characterise category structure as pairwise distances (e.g. using representational similarity analysis; RSA), which reveal whether categories are distinguishable, but not which representational axes organise the space as a whole. As a result, it remains unclear how low-level acoustic and higher-level semantic information are embedded as dimensions within the same functional structure of auditory cortex.

Here, we address these outstanding issues by deriving low-dimensional gradients from functional magnetic resonance imaging (fMRI) brain activity patterns (9). These gradients capture the major axes of functional organisation within human auditory cortex, and provide a common coordinate system in which all categories can be directly compared. This makes it possible to characterise how categories differ in their global organisational structure rather than in local activation patterns. We then project both an acoustic model space (based on gammatone filter-bank features) and a behavioural similarity space (derived from large-N human similarity judgements (10)) into this same gradient framework, allowing us to test how the brain’s representational axes align with low-level acoustic or higher-level semantic structure. Our high-depth dataset (three participants, 30 sound exemplars presented 56 time each, ∼4.7 hours fMRI acquisition each) ensures that these maps are robust and reproducible at the individual level, providing a strong basis for interpreting the cortical organisation of natural sound categories. The full fMRI dataset has been made publicly available (openneuro.org/datasets/ds006943).

## 2 Results

### 2.1 Task Design and Data Overview

Three participants completed 8 sessions of fMRI scans (56 runs total, 7 runs/session). In each run, participants heard all 30 sounds in a pseudo-random order, with jittered inter-stimulus intervals (8-17s) between each sound (see Section 4.3). To maintain attention throughout the runs, participants performed a 1-back task, pressing a button with the right index finger if they heard two sound exemplars in a row that belonged to the same category (Figure 1A). Per-session beta maps were estimated for each exemplar by concatenating runs within a session, yielding one beta map per sound exemplar per session (see Figure 1B and Sections 4.7). Gradients were derived by applying diffusion embedding (11) to the functional connectivity (FC) matrix of the fMRI time series (9). These beta maps were labeled by category to train Random Forest classifiers that learned both within-category similarity and between-category distinctions. Classifiers were trained and tested using cross-validation: 28 folds were generated by splitting sessions into 6-session training and 2-session testing sets, ensuring all runs from a session remained in the same set to prevent data leakage.

**Figure 1:**
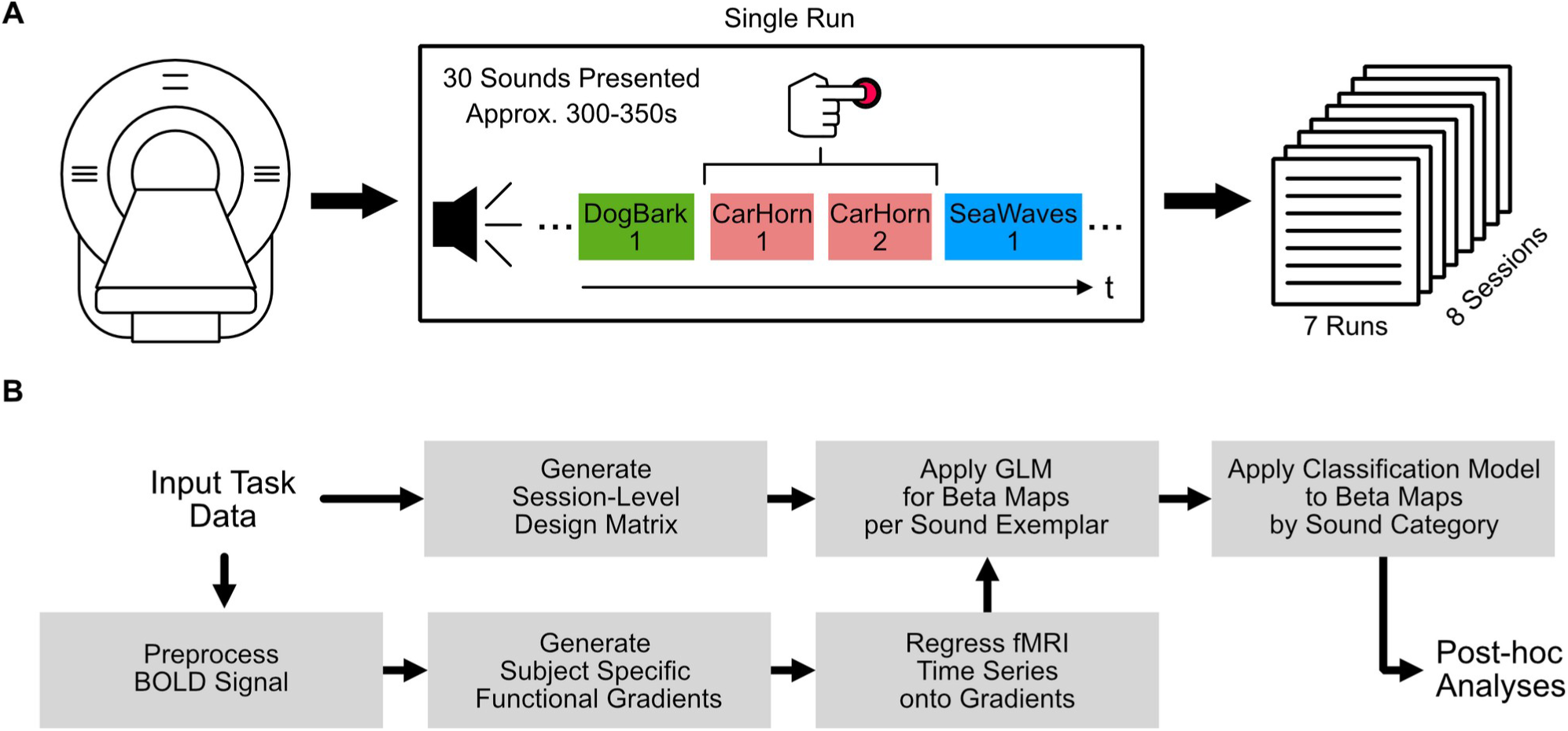
Schematic of fMRI Recording Task and Analysis Pipeline. **(A)** Schematic showing the task structure for each run during fMRI recording. While in the scanner, participants were exposed to 30 sound exemplars in a row. The sounds were ordered pseudo-randomly. Participants were instructed to press the button when they heard two sound exemplars that belonged to the same category. All 30 sound exemplars used in the study were played one time each every run, with order and ISI being the only variables that changed. Button press ordering was balanced across runs and sessions so every sound category had the same number of button presses overall. Participants performed this task for 7 runs in a session, and completed a total of 8 runs each. **(B)** Schematic showing pipeline of processing and analysis of recorded data. FMRI is preprocessed, and gradients are generated. The fMRI time series is then regressed onto these gradients to obtain one time series per gradient. The session-level design matrix is then applied to these gradient time series to obtain beta maps per sound exemplar from the gradient time series. These beta maps are applied to classification, which are then subject to further post-classification analyses.

### 2.2 Gradient-Based Representations Enhance Sound Category Classification in Auditory Cortex

As an initial step, we asked which cortical regions carried reliable information about sound categories, using two different cortical parcellations. *Csurf* auditory parcels (Figure 2A, top) delineate putative cortical areas on the basis of multiparameter maps, amplitude responses and tonotopic mapping (12). The *Schaefer*-1000 parcellation (Figure 2A, middle) segments the entire cortex into fine-grained functional regions (13). A third, hybrid parcellation (Figure 2A, bottom) combines the *csurf* auditory regions with anterior *Schaefer*-1000 regions and insula areas, creating a “frontal–auditory subset” to probe potential contributions from premotor and lateral prefrontal areas alongside auditory cortex.

**Figure 2:**
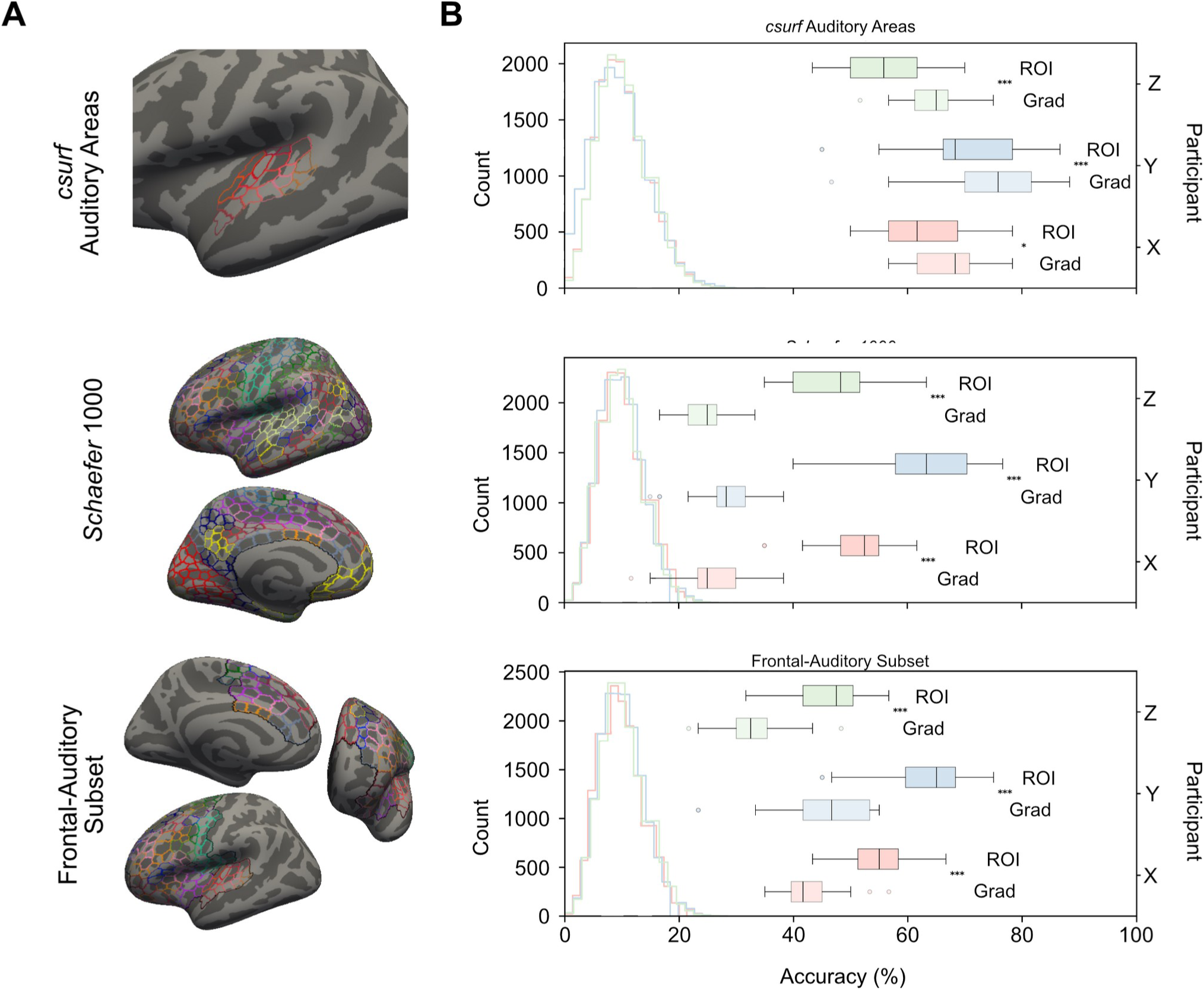
Cortical Parcellations and Classifier Accuracy of ROI and Gradient Embeddings. **(A)** Cortical parcellations used in both ROI and Gradient analyses. Left hemisphere of each parcellation projected onto an example subject surface in FreeView. **(B)** Accuracy of classification using the in-line cortical parcellation. Histogram shows the null distribution of classification accuracies from permutation tests across folds, with values corresponding to left y-axis. Box-plots show the observed accuracies across folds for each participant. Top box-plot of each participant is the accuracy for ROI classification, bottom is accuracy for gradient embedding classification. Colours denote participant. T-test results between ROI and gradient embedding approaches per parcellation per participant; * p<0.05, ** p<0.01, *** p<0.001.

We first used these parcellation schemes to provide regions of interest (ROIs) to classify the sound categories using a Random Forest classifier. In this ROI approach, beta coefficients were derived from mean values for each ROI, with the ROI beta values used as the features in the classifier. In Figure 2B, the box-plots indicate accuracy of the classifier accuracy for a given participant across 28 folds of the dataset. The top box-plot of each coloured pair indicates this distribution for the ROI approach. Each histogram shows the permutation tests results from 500 shuffles of category labels across folds, with each colour showing the null distribution of the given participant. Across participants when using ROIs, classification of the 10 natural sound categories (each with three exemplars) was highest using the *csurf* auditory regions (participant accuracy of 56-71% averaged across folds), demonstrating consistent and robust decoding of meaningful auditory objects directly from cortical activity. The *Schaefer*-1000 parcellation and frontal–auditory subset produced broadly comparable results (47-62% and 47-64%, respectively), with relatively minor inter-participant variability. Figure S1 shows that restricting the *Schaefer*-1000 parcellation to frontal areas alone reduced classification to within the tail end of the null distribution (20-33%), suggesting that classification primarily derives from auditory-related regions.

Feature importance reflects the percentage contribution of each variable to the model’s predictions. The auditory cortex emerged as the dominant region for predicting sound categories across all parcellation types in the ROI analyses, with feature importance values in the auditory cortex dominating those outside it (see Figure S2). This likely explains the superior performance of the auditory cortex in ROI analyses across participants, as the classifier primarily relied on category-related differences in this region, where other areas introduced information that would be considered noise for the discrimination.

We then applied the same Random Forest classifier approach to the gradient-derived data (Figure 2B, bottom-of-pair box-plots; see Section 4.10 for a description of gradient derivation and beta maps). Whereas in the ROI approach, each parcel is treated as an independent feature, in gradient embeddings, by contrast, beta maps can be interpreted as weights applied to region-spanning maps for each sound exemplar.

Categorisation accuracy for the gradient-based analysis, applied over the *Schaefer*-1000 and frontal–auditory subset parcellations the was significantly lower when compared to the more localist ROI analyses reported above; this held true for all participants comparing fold-wise distributions (Figure 2B).

By contrast, the gradient analysis over auditory cortex alone showed the ***opposite*** effect: accuracy significantly *increased* when applying gradient embedding, compared to the ROI approach. The fact that gradient beta-maps are represented via multiple cortex-spanning features rather than segregated ROIs likely explains why gradient embedding enhanced classification accuracy in the auditory cortex. We explore this phenomenon in greater depth in the next analyses.

### 2.3 Distributed and Category-Specific Organisation in Gradient Space

We further examined the auditory cortex gradient model to understand what drove its higher performance, relative to the parcel-wise beta maps approach. Confusion matrices and feature importance revealed that no single axis of the gradient space was sufficient for classification; instead, different sound categories recruited partially overlapping but distinct combinations of gradients. Figure 3 shows a cross-fold normalised analysis of gradient embedding results for the auditory cortex parcellation in an example participant. The fold-normalised confusion matrix (Figure 3A) indicates that certain natural sound categories—particularly Bird Song, Horse Galloping, and Sneezing—were consistently classified with higher accuracy.

**Figure 3:**
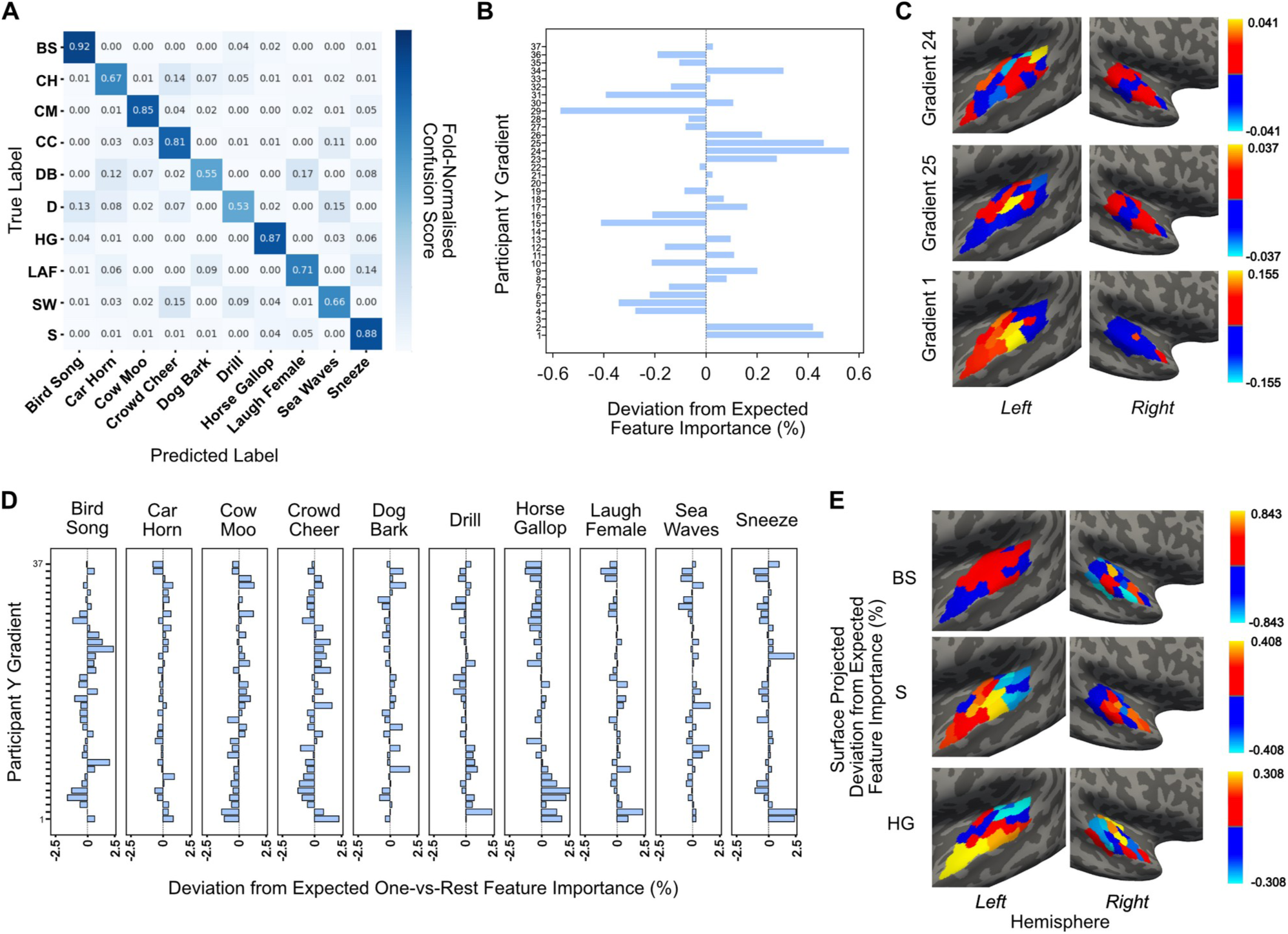
Example Participant Gradient Classification. **(A)** Fold-Normalised Confusion Matrix for example participant. Each row sums to 1, meaning each cell shows the percentage of time the given label on the y-axis was predicted as the label on the x-axis across folds. **(B)** Deviation of 10-category feature importance from expected uniform. Given 37 gradients, the expected importance is 2.70%, values given are percentage deviations from this value. **(C)** Cortical surface gradient maps of the most important gradients from B. Parcels are csurf auditory regions projected onto example participants surface. **(D)** Deviation from expected feature importance as in B, but for one-vs-rest classification. Each sub-plot is the feature importance deviation from 2.70% for predicting the given sub-plot’s category vs any other category (binary choice in classification). **(E)** Projection of gradient feature importance (from D) of three best performing categories in classification across folds (from A) onto the cortical surface. Each map is a weighted sum of all gradient maps, using corresponding feature importances as weights.

The matrix also indicates where the classifier more frequently confused particular category pairs across folds. For example, below the diagonal we observe a relatively frequent model confusion where the true label of Sea Waves was incorrectly predicted as Crowd Cheering 15% of the time (see Section 2.4 for details). This confusion was mirrored above the diagonal, where Crowd Cheering was predicted as Sea Waves 11% of the time. Category-pair confusions are not always symmetric: for example, Car Horn was predicted as Crowd Cheering 14% of the time in the example participant, but Crowd Cheering was only predicted as Car Horn 3% of the time. These patterns are indicative of underlying similarities between specific categories in the brain, which hint at auditory cortex organisational structure. Analogous confusion matrices for other participants are shown in Figure S3, and with similar cross-category confusions in the auditory cortex model.

Feature importance across the 37 gradients is displayed in Figure 3B for the example participant. Here, importance values are expressed relative to a uniform baseline (i.e., if all gradients contributed equally). Unlike ROI analyses, gradients do not capture the strength of activation in a given region. Instead, they instead represent axes of functional organisation as revealed by underlying brain dynamics. To illustrate the gradient approach to categorisation, we selected the gradients that were the most influential (24,25,1) and least important (29, 15, 31 and 5) for categorization.

Figure 3C shows the most influential gradient maps projected onto both hemispheres of Participant Y (see Figures S4, S5 and S6 for all gradients of all participants).

Each gradient has two opposing poles, meaning that areas at one extreme of the gradient are functionally opposed to those at the other. Gradient 24 shows these opposing poles in the medial–posterior area of the left hemisphere, with the putative caudo-lateral belt and A1 (most negative) and caudal parabelt (most positive) regions being at either end of the axis^1^. Gradient 25 distinguished the anterior-lateral belt from all other parcels, while Gradient 1 primarily separated anterior regions of the left hemisphere from the rest of the cortex.

Whilst these were the most important gradients in discriminating the categories, all gradients had varying degrees of importance when distinguishing the categories, with no one axis dominating in the category delineation, showing that multiple functional axes of information are required to distinguish sound categories. The organisation of gradients, and the most important gradients for each of the three participants, was not uniform. Figures S4, S5 and S6 show the first ten gradients of each participant. The location of poles of each gradient vary between participants significantly. This may in part be due to the bilateral nature of these gradients, where poles can span the auditory cortex of both hemispheres. It could also potentially be due to the parcellation approach used here. A voxel-wise approach could elucidate this further.

To examine how gradient importance varied by category, we used a one-vs-rest classification approach. In this procedure, the classifier was repeatedly trained and tested across folds, each time relabelling the data so that one sound category was classified against all others. Figure 3D shows the deviation from baseline feature importance of each gradient for each category. Again, these results indicate that no single gradient is sufficient: each category required a combination of multiple gradients for reliable discrimination. This further suggests that gradients act as axes along which different aspects of information are distributed, and that natural sound categories in the auditory cortex cannot be captured along a few simple dimensions of functional organisation.

To illustrate this, we projected the feature importance values onto the cortical surface for the three best-performing categories in the example participant (Figure 3E). Feature importance deviations (Figure 3D) were used as weights in a weighted sum of all gradient maps (see Equation 1 in Section 4.11). In these example projections, the Bird Song category showed relatively uniform contributions across the auditory cortex, with rostro-lateral areas (“RTL” and “R-A4”) contributing much less than the uniform baseline, indicating these regions contributed little information about this category. The Sneezing category relied on fewer gradients overall (Figure 3D), with anterior regions generally contributing more across both hemispheres, while posterior areas of the left hemisphere were less informative. For Horse Galloping, strong contributions arose from anterior temporal para-belt regions in the left hemisphere and from core and medial areas in the right hemisphere.

For clarity, these maps should not be considered “category maps”. Rather, they are visualisations of the combination of gradient axes that were utilised the most when delineating the given category from all others. These patterns suggest that different sound categories recruit partially distinct combinations of gradients, underscoring the distributed nature of auditory category representations. Despite each participant being represented by their own unique 37 dimensional space within which the sound categories were described by the classifier, the feature importance maps of each sound category show similarities between participants. (Cortical surface projections of feature importance for all categories and participants are provided in Figures S7, S8 and S9.) Whilst the intensity of specific regions within each feature importance map do vary between participants do vary, the major poles of importance/unimportance are consistent across participants. A visually clear example of this can be seen in the Sea Waves feature importance projection in Figures S7, S8, and S9 (though the pattern is evident in all categories). Although there is variation in specific intensities of importance between participants, the general pattern of importance in auditory cortices is consistent across participants for each of the category feature importance maps.

### 2.4 Bottom-up: Brain Representations Reflect Acoustic Drivers of Category Organisation

To understand how the classifiers utilised the functional axes descried above, and which gradients encoded them, we defined known axes of sound category discrimination, and embedded them in the gradient model We began with a well-established bottom-up representation of each sound exemplar by generating a gammatone representation for each stimulus (14), which is a cochlea-inspired decomposition of the sound into frequency channels. For each sound, we computed the mean envelope in each gammatone filter, yielding a compact frequency-based representation of the stimulus (Figure S10). Principal component analysis (PCA) across the 30 sound exemplars identified three components explaining 93.7% of variance (50.9%, 33.8%, and 8.9% for PCs 1 to 3), roughly corresponding to mid, low and high-frequency weightings (Figure 4A). We transformed each sound exemplar onto these axes, resulting in a 3-dimensional vector per sound exemplar. Using these vectors, we trained and tested classifiers equivalent to those trained on the fMRI data. With two cross-validation folds and 1000 repeats, the classifier achieved an average accuracy of 56.7%, while a permutation test with 500 permutations produced a null distribution with a tail at 20% (hence *p<0.002*). A comparison of the accuracy distribution across folds between the brain-based and gammatone models shows that the auditory cortex gradient representation captures more information about category structure (average accuracy 72%) than simple spectral properties (average accuracy 56.7%).

**Figure 4:**
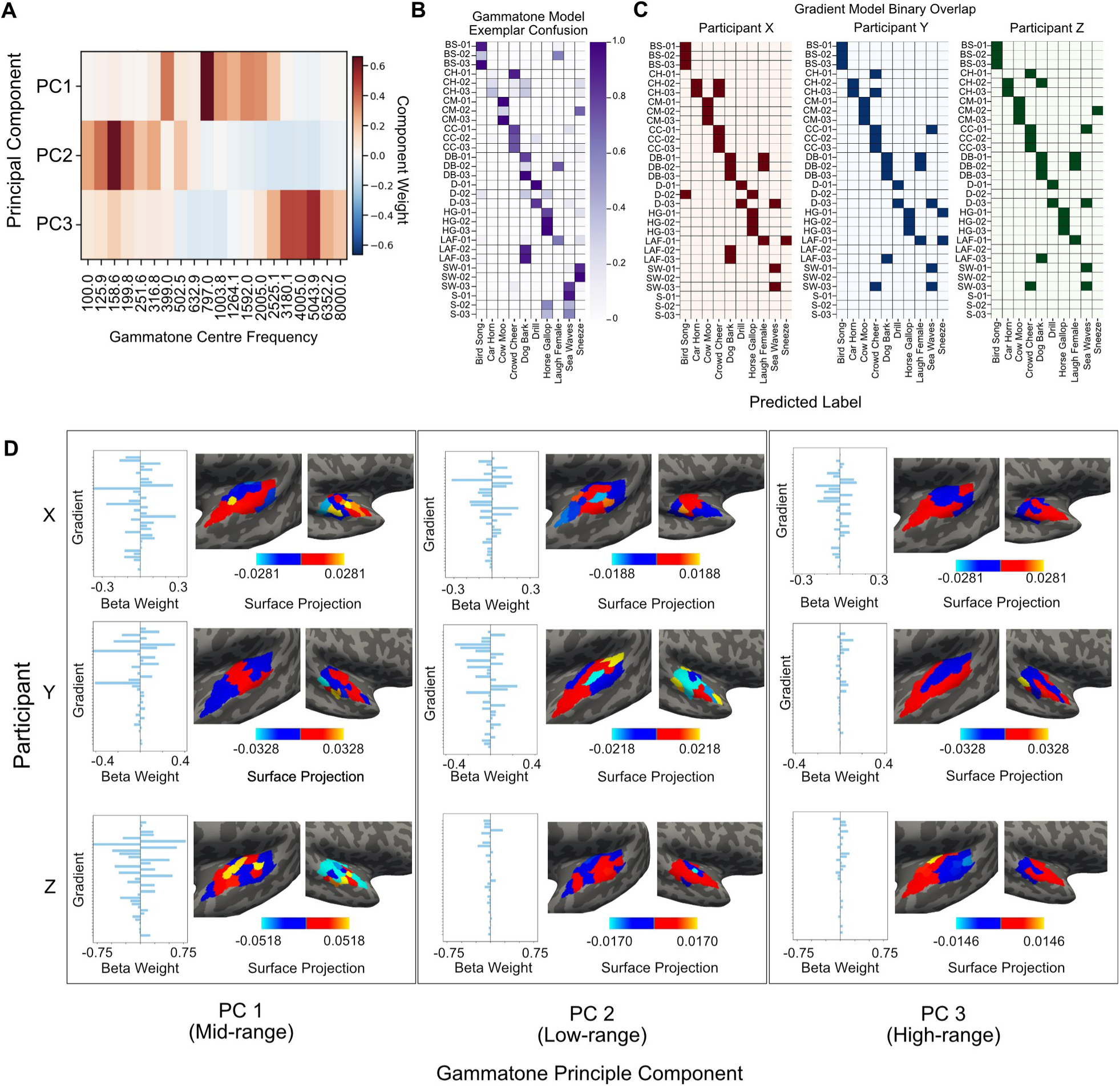
Comparison of Gammatone and Gradient Space Classifications of Sound Categories. **(A)** Principal Component eigenvectors of gammatone envelope means across all sound exemplars. X-axis shows the centre frequency of each filter. **(B)** Confusion matrix shows exemplar-level confusion matrix for classification using gammatone PC transforms of each sound category. Values are normalised so each row sums to 1, thus values are the percentage of instances the given exemplar was predicted as each category on the x-axis. (Note the “correct” diagonal is in vertical blocks of three). **(C)** Three confusion matrices are comparison between gammatone classifier performance (as in B) and each individual participant’s gradient-space classifier. If the same cell in both the gammatone confusion matrix and the gradient model’s confusion matrix is >0.1 (10%), that cell is highlighted in the binary overlap, indicating an overlap between the gradient and gammatone models. **(D)** Projection of gammatone-PC coordinates of each category (10 × 3) onto the category-specific gradient feature importance patterns (10 × 37; R^2^ values computed here). Each bar plot shows weightings of the projection of each PC, in each participant, onto the gradient space, with accompanying visualisation on the cortical surface using the gradient parcellation maps (38 × 37; Supplementary Figures 4, 5 and 6).

The permutation-normalised confusion matrix of exemplars for the gammatone classifier is given in Figure 4B. This plot is equivalent to Figure 3A, except that the “true label” axis now shows individual sound exemplars rather than their broader category labels. The gammatone classifier confusion matrix is followed by a binary overlap of confusion matrices between it and participant auditory cortex gradient models (Figure 4C). To illustrate the similarities between matrices, we highlight cells where both gammatone and auditory cortex gradient models predict that the exemplar is a given category > 10%. Misclassifications in the gammatone model are driven by exemplars with spectral profiles that closely resemble those of the categories they were incorrectly assigned to. It follows that when gradient models exhibit similar confusions, they are likely reflecting the same acoustic similarities between exemplars and categories, highlighting cases where category structure is predominantly shaped by acoustic features. Such examples can be found in Figure 4C; Car Horn is commonly mislabelled in both the gammatone and gradient models as Crowd Cheering, and Dog Barking often mislabelled as Laughing Adult Female. Additionally, the frequent confusions of the gradient model discussed in Section 2.3 (Figure 3A) - Sea Waves vs Crowd Cheering – are understood in Figure 4C as individual exemplars in each category that are acoustically similar to the other category. In two out of the three participants, the “Sea Waves 3” exemplar is mislabelled as Crowd Cheering with a frequency similar to the gammatone model, indicating that this exemplar specifically is likely driving the confusion between categories. The same goes the other way for “Crowd Cheering 1”. These specific exemplars likely overlap with the incorrect category pair in their spectral properties, causing the confusion in the auditory cortex model due to their similarity in their bottom-up representation of these specific exemplars.

To understand the way that the gradient space model may draw on spectral features, we embedded the gammatone representation of each sound category into the gradient space. To do this, for each category we averaged the principal component (PC) coordinates of each sound exemplar’s gammatone representation to obtain a single gammatone-PC category representation. Since the gammatone PCs and the fMRI gradient space share a category structure, we could then use the one-vs rest feature importance of each category as the target for a regression with the gammatone PCs (example shown in Figure 3D). This approach assumes that the acoustic PCs capture primary ways in which categories differ in sound space, and that classifier sensitivity to cortical gradients reflects, at least approximately, the same organisational structure of the categories.

A linear regression between the gammatone PCs and gradient feature importance yielded coefficient of determination (*R²*) values of 0.406, 0.377, and 0.367 for participants X, Y, and Z, respectively. This indicates that a substantial portion of gradient sensitivity to sound categories is explained by low-level acoustic features, though the moderate *R²* values suggest that other factors also contribute. Figure 4D shows each of the three gammatone PCs weightings when projected into the gradient space of each participant (bar plots). These weightings have in turn been used for visualisation by projecting the weights onto participants’ cortical surfaces (surface projections; see Equation 1 in Section 4.11). This allows us to see how axes of acoustic variation are expressed across gradients, rather than mapping categories to discrete regions. These projections vary across participants, which may partly reflect differences in classifier performance as well as underlying tonotopic map topography. However, as Figure 4D illustrates, participants also differ in the degree to which their gradient models reproduce exemplar-level gammatone confusions. This suggests that individual variability in how acoustic structure shapes category organisation could underlie some of the observed differences in gradient–auditory-PC alignment.

### 2.5 Top-Down: Behavioural Judgements are Reflected in Brain Category Structure (but Less than Acoustics)

After confirming overlap between acoustic information and brain category structure using gammatones, we next investigated whether top-down, behaviourally-driven distinctions also aligned with cortical representations. To do this, we used data from a previous behavioural study in which participants organised the same sound categories according to their perceived semantic dissimilarity (15). A representational dissimilarity matrix (RDM; Figure 5A) from that study was subjected to PCA, extracting three principal components to capture the main axes of perceived category differences, effectively establishing a behavioural-judgement-based category structure. (The three PCs explain 78.8% of the variance; 46.6%, 22.5%, and 9.6% for PCs 1–3) The placement of each sound category along these three behavioural PCs is shown in Figure 5B.

**Figure 5:**
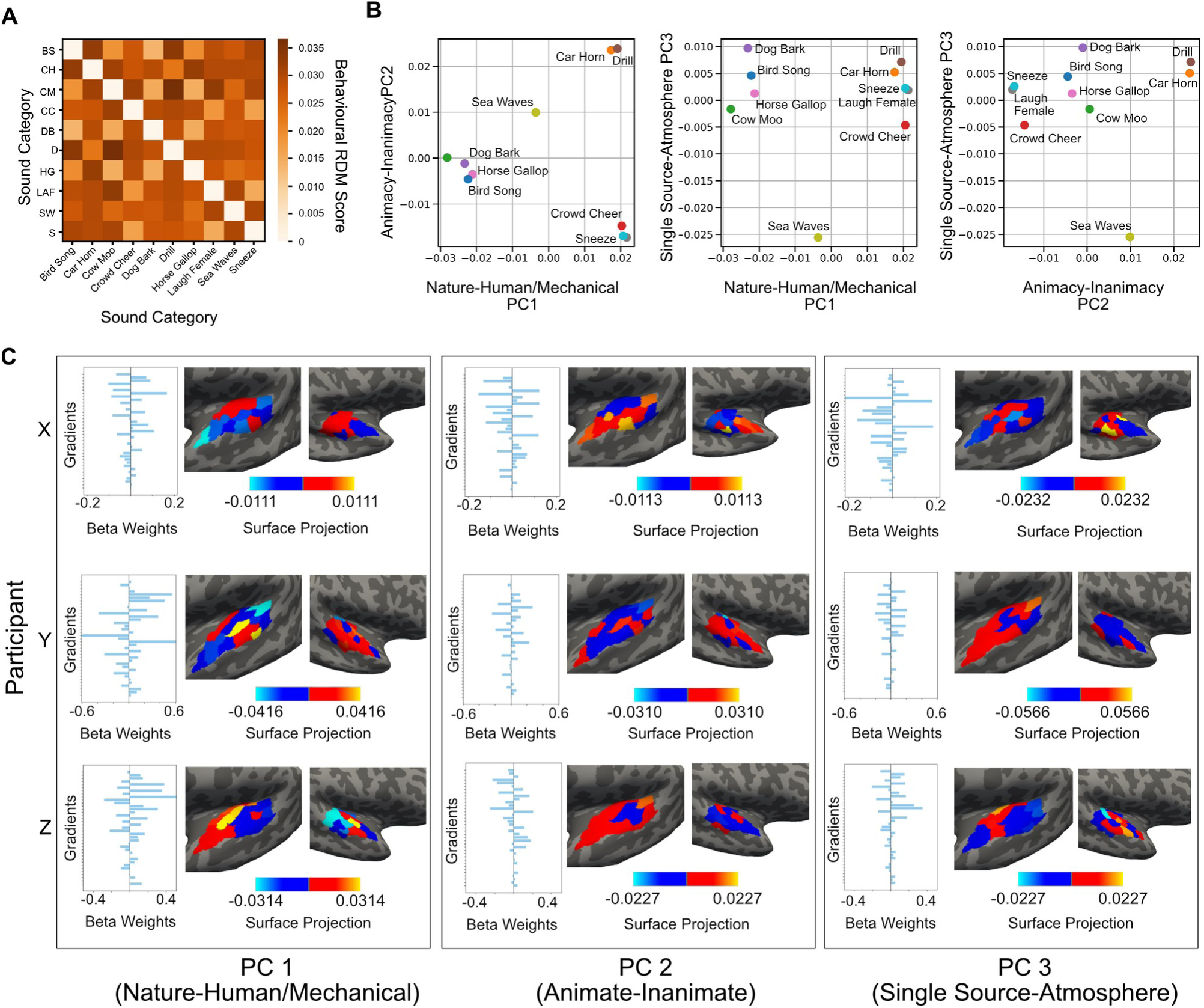
Semantic Behavioural Judgement Projected into Functional Gradient Space. **(A)** Representational Dissimilarity Matrix of sound categories from Semantic Behavioural Judgement study using our sound categories. **(B)** Sound categories in the behavioural RDM space after PC transform. Three PCs were fitted to the RDM, and categories were transformed onto those three axes. Each sub-plot shows a intersecting plane of the three axes. Colours indicate sound category. **(C)** Projection of behavioural PC axes onto each participants’ gradient axes. Bar plots show the weighting of the given behavioural PC onto all 37 gradients of the given participant. These weightings were then projected onto participant surfaces. Each row shows a participant, each column the PC. The same behavioural PCs were projected onto all participant surfaces through their own gradient classifier. PCs have been labelled to show a simple negative-positive axis of potential semantic meaning.

PC1 primarily separates naturally occurring sounds from human-generated or technologically-mediated ones. Negative loadings correspond to sounds produced by animals or the natural environment (e.g., Cow Moo, Bird Song, Horse Galloping, Sea Waves), while positive loadings correspond to sounds originating from human actions or machinery (e.g., Sneezing, Drilling). PC2 appears to capture animacy: positive values correspond to inanimate sources (Drilling, Car Horn, Sea Waves), whereas animal- and human-sourced sounds have negative values, with humans showing the strongest negative loadings. PC3 is less interpretable but may reflect sounds associated with a larger environment or atmosphere as a source: SeaWaves and CrowdCheering have negative values, while other sounds that have a “single source” are near zero or weakly positive.

We used these behavioural axes in a linear regression with the gradient feature importance of each category as the target (Figure 5C, bar plots), analogous to the approach applied to the gammatone PCs in Section 2.4. This analysis yielded *R²* values of 0.321, 0.296, and 0.297 for participants X, Y, and Z, respectively, demonstrating that auditory cortex representations are meaningfully shaped by behaviourally judged semantic distinctions. While these values are slightly lower than those obtained for the gammatone axes, they nonetheless indicate that top-down, semantic information contributes substantially to the organisation of category representations in the auditory cortex. This effect is consistent across participants, and suggests that category structure in auditory cortex is influenced by both bottom-up acoustic features and top-down semantic information. The difference in the predictive power of the gammatone- and behaviourally-derived PCs indicates that these two sources of information contribute uniquely to shaping cortical category representations. The mapping of behavioural PCs onto the gradient space was projected onto each participants’ surface for visualisation (Figure 5C). These can be conceptualised as the semantic dimensions within the gradient space along which the categories are distinguished in each participant. Similarly to the gammatone approach, the weights and maps are not consistent across participants.

### 2.6 Flexible Representation of Category Structure in the Brain

To investigate how the influence of the gammatone and behavioural axes on the gradient classification model may overlap or differ, we investigated the category structure within each space and compared them. To do this, we took a representational dissimilarity matrix (RDM) from the three spaces; gammatone, behavioural, and gradient classification. Figure 6A presents three RDMs that show the differences between each sound category in three different contexts (for participant Y). In the functional gradient space RDM, each cell’s shading shows the dissimilarity (in Euclidean distance) between the feature importances for a pair of sound categories, where feature importance is defined as the classifier’s weights on each gradient axis from the One-vs-Rest model (see Sections 4.8 and 4.10). The gammatone and behavioural RDMs show the Euclidean distance between the PC eigenvalues of each pair of categories in the gammatone and behavioural spaces.

**Figure 6:**
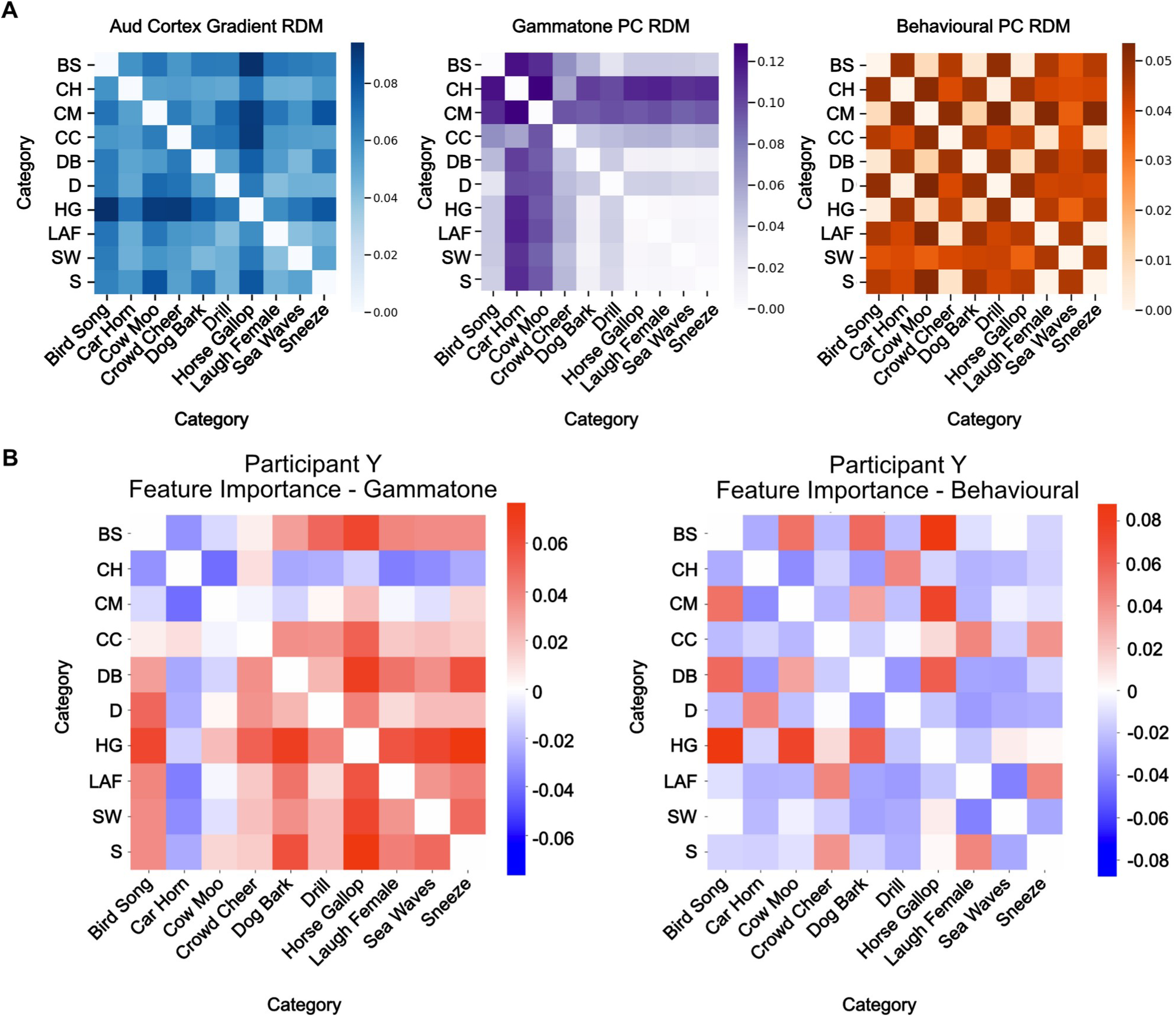
Comparison of Category Structure between Brain Data Classification with Gammatone and Behavioural Projections. (A) Distance Matrices between categories in each of the representational spaces. From left to right, participant Y feature importance projection distances, gammatone PC projection distances, behavioural PC projection distances. (B) Comparison of pairwise distances between category pairs for example participant Y feature importances, and behavioural (left) and gammatone (right) PC projections. Points are given as colour pairs to show each category pair. Midline indicates where x and y axis are equal.

These distances represent category distinction in each of the spaces – a larger distance between two categories in the gammatone model would indicate that they are more spectrally distinct, and a larger distance in the behavioural model would indicate that they are deemed more distinct in a general semantic judgement. To understand to what extent these acoustic and semantic category structures are similar or different to the gradient representation, we computed the difference between the RDM of the brain model, and the RDM of both the gammatone and behavioural models. Figure 6B shows the difference between the RDMs of the brain model and the gammatone (left) and behavioural (right) RDMs in example participant Y. Positive values indicate that a category pair is more dissimilar in the brain model, and negative values indicate the category pair is more dissimilar in the gammatone (left) or behavioural (right) models. If the value is zero or near zero, it indicates that the brain model category structure is the same or very similar to the gammatone or behavioural model.Whilst general category structure in the brain model very rarely showed a strong similarity to either the gammatone model or the behaviourally derived model, some category pairs were comparable. For example, Bird Song and Sea Waves showed a larger difference in the brain model than in the gammatone model, indicating that the brain model distinguished between these categories with more than just spectral information. When reviewing this same category pair in the comparison of brain and behavioural models, we see that the difference is effectively zero. This suggests that this part of the category structure is mediated primarily by semantic information in the brain, rather than acoustic information. Similar observations can be made for the Horse Galloping vs Sneezing, and Drilling vs Crowd Cheering pairs. The opposite trend is also observed: the dissimilarity of the Drilling vs Cow Moo category pair in the brain was very similar to the gammatone model, but different to the behavioural model. This indicates that this part of the category structure is mediated moreso by acoustic information than semantic information.

Some category structure in the brain also appears to not be entirely described by just the gammatone or just the behavioural model. For example, Horse Galloping and Bird Song are strongly dissimilar to one another in the brain representation – much more than they are distinguished in the gammatone model or behavioural model. Additionally, some categories do appear to be more acoustically distinct from others. Car Horn and Cow Moo both show larger distances from other categories (Figure 6A, middle). Accordingly, when comparing feature importance to gammatone distances (Figure 6B, left), these categories typically show greater distances from other categories in the gammatone model than in the brain model. Hence this pattern is likely due to spectral the distinctiveness of these categories.

The equivalent plot to Figure 6B is given for all participants in Figure S11, with an average across participants also shown. Remarkably, the general category structure shown in Figure 6B is very consistent across participants. This indicates that, although the spatial arrangement of gammatone and behavioural axes varies between participants (Figure 4D and Figure 5C), the pairwise representational geometry of sound categories within auditory cortex is highly consistent across individuals.

These observations suggest that the brain may utilise different types of information to distinguish between categories. Some category pairs show greater alignment with acoustic differences, as captured by the gammatone comparison (Crowd Cheering vs Sneezing), while others align more closely with behavioural or semantic distinctions (Drilling vs Laughing Adult Female). For some pairs, both sources distinguish categories more readily than the brain model (Car Horn vs Cow Moo), and for others, the brain representation delineates them more than both sources (Dog Bark vs Bird Song). Overall, this is consistent with the idea that the brain flexibly weights different features when representing category structure.

## 3 Discussion

Methodologically, our finding that gradient-based models within auditory cortex explain category structure better than ROI-based models extends gradient and multi-scale frameworks—developed largely for resting-state connectivity and low-level topographies—into the domain of task-evoked natural sound categorisation. Approaches such as (9) have shown that cortical organisation can be captured as smooth gradients. Prior work on sound category representations has typically characterised them using ROI- or searchlight-based approaches (3–5,16,17). Our results show that modelling auditory cortex using gradients yields measurably superior accounts of category geometry, indicating that category-relevant information is organised along functional axes that ROI approaches partially obscure.

Notably, the advantage of gradient-based modelling was specific to auditory cortex: extending models to frontal or whole-brain voxels did not yield comparable gains. This pattern aligns with evidence that reliable natural-sound category geometry is strongest in superior temporal cortex (4,5,17), whereas frontal contributions are more task- and state-dependent (7), diluting stimulus-tied structure when aggregated at whole-brain scale. This result also aligns with known risks that high-dimensional whole-brain feature spaces can obscure signal without strong inductive biases (18). Future studies could build on these findings by examining voxel-level organisation within auditory cortex to capture finer-grained representational structure and utilise approaches that extend beyond classical auditory cortex without losing predictive power.

Category representations were distributed across the full set of gradients rather than being explained by any single organisational axis. This suggests that the dominant functional axes of activation in auditory cortex—which are thought to reflect low-level acoustic features such as tonotopy and spectral–temporal tuning (19–21)—were not themselves the dominant axes of category structure, even though they contributed to category decoding. Rather, category structure appeared to be embedded across multiple axes, consistent with prior findings that category distinctions are superimposed on pre-existing acoustic organisation (4,16). To assess whether the functional gradients simply recapitulated tonotopic organisation, we compared each participant’s tonotopy map to their 37 task-derived gradients (See Figure S12). All gradients correlated with the tonotopy map to varying degrees, and together explained nearly all variance in tonotopy (*R²* ≈ 0.99), but no single gradient aligned strongly with it. This indicates that the gradients recover tonotopic information as part of a broader representational space, rather than reflecting tonotopy alone. Accordingly, category structure was distributed across gradients that combined tonotopic and non-tonotopic information, rather than being aligned with the primary frequency axis.

We found that feature-importance patterns derived from fMRI activity in auditory cortex captured category structure more effectively than models based solely on principal components of gammatone filter-bank features, which approximate early cochlear processing (21), indicating that gradients incorporate additional non-spectral dimensions. This aligns with prior findings that higher-level categorical structure emerges in non-primary auditory regions beyond what can be predicted from low-level acoustic features (23,24).

We then examined how acoustic and semantic structure contribute to the organisation of sound categories in auditory cortex by projecting gammatone- and behaviour-derived representational axes into the brain gradient space. To our knowledge, this framework of a shared representational space is the first of its kind. With the brain model as the target, we found that the gammatone axes explained a substantial proportion of variance (average *R²* = 0.383), consistent with previous reports that spectro-temporal features account for much of the representational structure of primary and early auditory cortex (5,16). Behaviour-derived axes explained a lower but still meaningful proportion of variance (average *R²* = 0.305). However, because behavioural similarity judgments often incorporate acoustic regularities, we cannot interpret this *R²* as reflecting purely semantic structure. Instead, the neural predictability of the behavioural model likely reflects a mixture of conceptual structure and acoustically mediated similarity that naturally co-occur in many real-world sound categories. This interpretation aligns with prior evidence that both acoustic and higher-level semantic attributes shape representations in non-primary auditory cortex (6,7). Nonetheless, the fact that the behavioural model explains reliably less variance than the gammatone model suggests that acoustic structure exerts a stronger influence on cortical organisation than the semantic associations captured in human judgments, even when accounting for the possibility of shared acoustic–semantic variance. Both sets of behavioural axes were weighted variably across multiple gradient axes, with no single gradient functional axis aligning totally with any single gammatone or behavioural axis. These findings indicate that both low-level spectral and higher-level semantic dimensions are reflected within the representational geometry of auditory cortex, with acoustic information exerting a somewhat stronger influence. Moreover, the difference in predictive power between the two model spaces suggests that they capture partially distinct aspects of cortical category structure rather than reflecting a single underlying representational dimension.

To probe how acoustic and semantic information contribute to category structure, we compared the geometric arrangement of categories across brain, gammatone, and behavioural spaces using Representational Dissimilarity Matrices (RDMs). Across category pairs, the brain’s representational geometry reflected a variable mixture of acoustic and semantic structure: some pairs were organised primarily by spectro-temporal similarity, others by higher-order distinctions, and many by both. Although each participant’s gradients and axis topographies were idiosyncratic, their projected category geometries converged on a highly similar structure. In other words, participants used different spatial representations to realise the same underlying representational geometry of natural sound categories. This pairwise heterogeneity is expected if auditory cortex contains intermediate, non-spectral population axes, and if primary vs. non-primary fields differ in their balance of acoustic and higher-order structure (24–26). It is also consistent with evidence that different areas of auditory cortex differentially contribute acoustic and non-acoustic information (27). In this model, our gradients provide a unified representational space in which these known contributors are embedded as part of a global functional geometry. Mechanistically, it has been shown that rodent ACx exhibits discrete, attractor-like population modes whose mixtures are biased by learning, promoting generalisation among already-similar sounds (28–30). This offers a plausible circuit-level basis for the heterogeneous weighting we observe across category pairs.

Several limitations of this study should be considered. Although the sound categories used here were diverse, they capture only a subset of the richness of real-world auditory experiences. Use of a wider category space could benefit investigation of the questions addressed here. Additionally, the use of cortical parcellations, while improving interpretability, also introduces spatial coarseness that may obscure fine-grained category representations within auditory cortex. Future study should consider voxel-level analysis using the approaches applied here. Further, comparison of the gradient model to state-of-the-art DNN models (23,26,31–33) may confirm the robustness of the approach. Finally, our models were based on task-evoked responses, which might differ from the brain’s intrinsic functional organisation; comparing task-evoked gradients with resting-state gradients could clarify how stable these representational axes are and how they support perception and cognition.

Overall, our findings suggest that natural sound categorisation is supported by distributed, multi-dimensional representations in auditory cortex, where category structure emerges from the integration of both low-level acoustic and higher-level semantic information. This flexible representational scheme may enable the auditory system to adaptively prioritise different types of information depending on context and task demands, supporting robust and generalisable sound recognition in the complex environments we encounter every day.

## 4 Methods

### 4.1 Participants

There were 3 participants in the study. All participants were female, right-handed, and within the age range of 30-40 years. None reported any hearing impairments or neurological conditions. All participants provided written informed consent prior to the experiment, and the study was approved by the local ethics committee at University College London.

### 4.2 Natural Sound Exemplars

Ten natural sound categories were chosen from a selection of 53 sound categories in an existing natural sounds dataset (10). The ten natural sound categories used were Bird Song, Car Horn, Cow Moo, Crowd Cheering, Dog Barking, Drilling, Horse Galloping, Laughing Adult Female, Sea Waves, and Sneezing. All sounds lasted 2-3 seconds. Three exemplar sounds were used per category, which were chosen from a selection of 10 exemplars per category. The three chosen were those with the highest ranking of sound category representativeness (34).

### 4.3 Experimental Design

Eight separate sessions of recording were carried out per participant over the course of three months, with each recording session consisting of 7 runs of the experiment. Each run lasted on average 296.57 seconds (range *281-390s*, *σ=8.87s*, total of all runs approx. 4.7 hours per participant).

For each run, each sound exemplar was played a single time. Participants engaged in a one-back task, instructed to press a button with their right index finger when two sound exemplars from the same category occurred sequentially in the sequence of sounds heard. There were four button press events per run (the participants were not made aware of this).

For each of the seven runs of a session, both the order of sound exemplars and the inter-stimulus interval (ISI) were pseudo-random. A simple shuffling algorithm was used to generate the order of sound exemplars for the seven runs in a session. The 30 sounds were shuffled multiple times to find valid and unique sound orders for a run. A shuffle was used as a run’s sound order if and only if the shuffle contained four instances where two exemplars of the same category were next to each other in the order of sounds. This process of order generation was carried out for every run in a session. Sessions were then checked manually to ensure that there were no unusual sequences (for example many runs with 3 button presses in a row, or runs with many button presses for the same category), and were edited accordingly. Manual editing of generated sequences ensured that the number of button press events per sound category was uniform across the 8 sessions.

The ISI between sounds was chosen by sampling from a probability density function (PDF). A truncated normal PDF was used (*μ = 9.0s*, *σ = 1.5s*) with lower and upper bounds of *7.0s* and *18.0s* respectively. The PDF was sampled 30 times per run with each sample being used as the duration of silence proceeding each sound exemplar. The same run-level sound orders were used for each participant, but the order in which the runs were exposed was shuffled between participants.

### 4.4 FMRI Data Capture

Images were acquired with a 3T Siemens Prisma scanner and 32-channel head coil (Birkbeck-UCL Centre for Neuroimaging). Head movement was minimised with a head stabilisation prototype (MR Minimal Motion System). Audio was delivered to participants over Sensimetrics earbuds with pre-filtering to accommodate the earbuds’ response profile.

Structural images were acquired using a T1-weighted magnetisation-prepared rapid acquisition gradient echo (MPRAGE) sequence (*TR = 1.53s*, *TE = 3ms*, *FOV = 256 mm*, *flip angle = 9°*) with 1mm sagittal slices. Functional echo planar images (EPI) were acquired using a T2*-weighted sequence (*TR = 1.0s*, *TE = 1.0s*, 2mm thickness, 48 slices, in-plane resolution *= 2mm × 2mm*, *FOV = 106mm*, flip angle *= 60°*, multi-band acceleration factor *= 4*). Eight dummy volumes were included at the start of each scan to allow the scanner to reach B1 equilibrium. Functional images were acquired with anterior-to-posterior phase encoding. Following each run two volumes with reversed phase encoding were collected to correct for image distortion caused by B0 inhomogeneities.

### 4.5 Preprocessing

FMRI processing was carried out using the *FSL* preprocessing pipeline. The following preprocessing was applied at the run-level. *Top-up* was applied to apply phase encoding corrections (35), the output of which was applied to *FEAT* Version 6.00. Registration from EPI to high resolution T1 structural images was carried out using *FLIRT* (36). Registration from high resolution structural to standard space was then further refined using *FNIRT* (37). The following pre-statistics processing was then applied; motion correction using *MCFLIRT*; non-brain removal using *BET* (38); spatial smoothing using a Gaussian kernel of full-width at half maximum (FWHM) *2mm*; grand-mean intensity normalisation of the entire 4D dataset by a single multiplicative factor; high pass temporal filtering (Gaussian-weighted least-squares straight line fitting, with *σ = 50s*). Independent Component Analysis (ICA)-based exploratory data analysis was carried out using *MELODIC* (39), in order to investigate the possible presence of unexpected artefacts or activation. Automatic classification and removal of noise Independent Components (ICs) was then carried out using *ICA-AROMA* (40), which utilised the warp transforms generated from *FNIRT*. Prior to analysis, voxel-wise standardisation was performed using the *RobustScaler* from *scikit-learn* (*41*), which removes the median and scales the data according to the interquartile range, in order to control for inter-run and inter-session variability.

### 4.6 FMRI Parcellation Masks

Native space cortical surfaces for each participant were generated using *FreeSurfer’s* structural processing stream (42). The Schaeffer-1000 cortex parcellation (13) and the auditory regions of the *csurf* tonotopic parcellation (12) were used, along with a combination of the two. The combination mask included the precentral sulcus and insula regions of the Schaefer-1000 parcellation, minus medial frontal pole areas. These frontal areas were masked with the auditory regions of the *csurf* mask on the *fsaverage* surface. In any overlap of vertices between the two masks, the *csurf* label took priority. Each parcellation was projected from *fsaverage* space onto each participant’s cortical surface in turn. The native volumetric space of each run was registered to each participant’s cortical surface using *bbregister* (*42*). The generated transform was then used to project both parcellations onto each run as a mask. The mean value of all voxels within each parcel at each time point was calculated to generate a parcellated time series for each mask.

### 4.7 ROI Beta Coefficient Maps of Sound Exemplars

Timing regressors were generated using *nilearn*’s first-level design matrix functionality (43), convolved with the Glover haemodynamic response function (HRF; (44)) as implemented in *nilearn*, using the default parameters (temporal oversampling=50, response length=32s, onset=0s). Linear regression was then applied to the concatenated BOLD signal from each session using one convolved regressor per sound exemplar, producing a beta coefficient map for each exemplar.

These beta maps were computed across the seven concatenated runs within each session, resulting in a single beta map per exemplar per session. To account for low-frequency fluctuations and baseline shifts at the run level, a third-order polynomial drift model and constant term were included, yielding four nuisance regressors per run. Button press instances were also used as nuisance regressors. Thus, each session produced one beta coefficient per sound exemplar. With eight sessions per participant and three exemplars per sound category, this resulted in 24 beta coefficients per sound category (3 exemplars × 8 sessions).

### 4.8 Classification of Sound Categories

Each of the 24 beta maps per sound category (3 exemplars × 8 sessions) was assigned the corresponding sound category label. To classify sound categories, the *scikit-learn* implementation of the Random Forest classifier (41) was trained and tested using these beta maps. Importantly, data were split by session: beta maps from 6 sessions were used for training, and the remaining 2 sessions were used for testing. This was done to prevent overfitting to session-specific noise and to promote generalisability. All 28 possible 6–2 train–test splits of the 8 sessions were evaluated using cross-validation. Classification accuracy was recorded for each fold.

To assess statistical significance, a permutation test was performed for each fold. The training labels were randomly shuffled 500 times, and a new classifier was trained and evaluated for each shuffle. The distribution of accuracies from the shuffled data was compared to the accuracy from the real labels. A *p*-value was computed as the proportion of shuffled accuracies that exceeded the real accuracy; i.e., if no shuffled accuracy exceeded the real accuracy, the resulting *p*-value would be *0.002 (1/500)*. The average accuracy across all 28 cross-validation folds was taken as the overall accuracy of the model. A Random Forest classifier was subsequently trained on all data for further scrutiny.

For each fold, the feature importance was computed, providing a measure of how strongly each feature contributed to distinguishing between sound categories. Feature importance in Random Forests is derived from the reduction in impurity (Gini impurity) that results from splitting on a given feature across all trees in the ensemble (45). Features with higher importance scores indicate features that consistently contributed to improving category classification. The feature importance was computed on all folds and averaged across folds, for each participant. Importance values within a single model sum to 1.

### 4.9 One-vs-Rest Classification

One-vs-rest (OvR) classification is a strategy for handling multi-class problems using binary classifiers. Rather than separating all classes at once (as in Section 4.8), the approach decomposes the problem into multiple binary tasks. For each class *c*, a separate classifier is trained to distinguish that class (the “one”) from all other classes combined (the “rest”). If there are *C* classes for a given fold, then *C* binary classifiers are trained for that fold. The *c^th^* classifier is trained with label 1 if it belongs to the given class, and 0 if it does not. For each fold, a new sample is passed through all *C* classifiers. Each classifier outputs scores equivalent to those laid out in Section 4.8, only for the given category vs all others.

### 4.10 Gradient Analysis

To compute functional gradients for each parcellation, we used the *BrainSpace* toolbox (46). For each training fold, we computed the pairwise Pearson correlation coefficient between the BOLD time series of all parcels. For each fold of the data (as described in Section 4.8) the preprocessed fMRI was used to produce a symmetric Functional Connectivity (FC) matrix. To reduce the dimensionality of these FC matrices while preserving their geometric structure, diffusion embedding (11) was applied to generate 60 gradients per fold in the Schaefer and combination parcellations, and 37 gradients in the auditory cortex parcellation (38 parcels).

A low-dimensional representation of each session-level beta coefficient map was generated by regressing each gradient component onto the BOLD time series of each session, yielding a time series for each gradient. We then regressed the design matrix (described in Section 4.7) onto these gradient time series to obtain beta coefficients representing each sound exemplar embedded in gradient space. The classification laid out in Section 4.8 was subsequently applied to these embedded beta maps for each parcellation. Note that gradients were computed separately for each fold and used exclusively within that fold. For visualisation, the gradients of each fold were aligned via Procrustes rotation, taking the first fold as reference. The mean across rotated gradients is shown in visualisations. Similarity of the rotated gradients was computed across folds to ensure consistency.

### 4.11 Feature Importance Projections

To represent the feature importance output of gradient classifiers on the surface, we used feature importances as weights for each gradient map (derived as described in Section 4.10). Equation 1 shows the weighted sum used to compute this mapping:

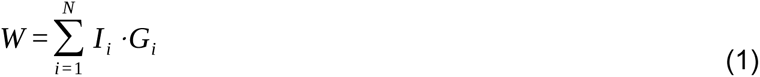

where *W* is the weighted map, *N* is the number of gradients, *I* is the feature importance for the given gradient and classifier, and *G* is the set of gradients. This approach was applied for outputs of both the general classifier models with 10 categories (as described in Section 4.8), and each of the One-vs-Rest models (as described in Section 4.9), generating a weighted map that distinguishes one sound category from all others.

### 4.12 Behavioural Data

As described in (34) 73 participants arranged a subset of the sounds from the original set of 53 categories in a circular arena based on their understanding of “meaning-based similarity” of the sounds heard. The sound exemplars heard were those rated with the highest “category representativeness”. Participants were fluent English speakers, and participants that failed volume checks or showed random patterns of responses or unstable arrangements were excluded from the analysis. Pairwise distances between categories were measured as Euclidean distance. The subset of 10 categories used in this study were isolated from that study (Figure 5A), and used as the basis for behavioural axes representations (Figure 5B).

### 4.13 Projection of Gamma-tone and Behavioural Data to Functional Gradient Space

To examine the correspondence between spectral properties of the sounds and auditory cortex gradient organisation, we first extracted gamma-tone features from each exemplar, averaging across exemplars within each sound category to yield a category-by-frequency-band representation (10 categories × 20 frequency bands). We then applied principal component analysis (PCA) with three components to this matrix to identify the dominant axes of variation in the spectral profiles across frequency bands. The PCA loadings defined frequency-band–by–principal component mappings, which we used to project each sound category into a low-dimensional space, resulting in a 10 × 3 matrix of category scores along three principal components. In parallel, category-specific feature importances were estimated using a one-vs-rest classification of sound categories (one category vs all other categories). These feature importance values were averaged across cross-validation folds to yield a gradients-by-categories matrix (37 × 10). Because both representations share the same set of sound categories, we fit a linear regression between the gradient-based feature importance values and the category scores in gamma-tone—PC space. This regression produced a gradients-by-PCs projection, allowing us to assess the degree to which cortical gradient organisation aligned with the low-dimensional spectral axes of the sound categories. Model fit was quantified using the coefficient of determination (*R²*).

We repeated this procedure using behavioural data obtained from a semantic judgment task. Specifically, we constructed a representational dissimilarity matrix (RDM) across the 10 sound categories based on participants’ similarity judgments. Principal component analysis was then applied to the RDM to extract the dominant dimensions of semantic variation, and each category was projected into this space, yielding a 10 × 3 category-by-PC representation. As with the acoustic features, we aligned these behavioural PCs with cortical gradient feature importance values by fitting a linear regression between the one-vs-rest feature importance matrix (37 × 10) and the behavioural PC scores (10 × 3). This produced a gradients-by-PCs projection reflecting how semantic similarities between categories were expressed in gradient space. Model fit was again quantified using *R²*, allowing us to compare the degree to which cortical gradients captured acoustic versus behavioural dimensions of category organisation. Visualisation of PCs on the cortical surface used the weighted mapping approach laid out in Section 4.11 and Equation 1.

Mapping the PCA axes of both the acoustic and behavioural spaces onto the gradient-based feature importance patterns assumes that these axes capture the primary ways in which categories differ within each domain, and that the classifier’s sensitivity to cortical gradients reflects, at least approximately, these same organising dimensions.

## Statistical information

All statistical analyses were conducted in Python using the *scikit-learn* library for classification and *numpy*/*scipy* for statistical calculations. The primary analysis used a Random Forest classifier to predict sound categories from fMRI beta maps. For each participant, 24 beta maps were obtained per category (3 exemplars × 8 sessions), resulting in *n* = 240 beta maps (10 categories × 24 beta maps per category).

Classification performance was evaluated using an exhaustive cross-validation scheme in which data were split by session to avoid overfitting to session-specific noise. In each fold, beta maps from six sessions (*n* = 180 maps) were used for training, and beta maps from the remaining two sessions (*n* = 60 maps) were used for testing. All 28 possible 6–2 train–test splits of the 8 sessions were tested, yielding 28 accuracy values per participant.

Statistical significance was assessed separately for each fold using a permutation test with 500 label shuffles (*n* = 500 permutations). For each shuffle, the classifier was retrained and evaluated on the same train–test split to generate a null distribution of accuracies. The *p*-value for each fold was computed as the proportion of permuted accuracies exceeding the true accuracy. Because no permutation accuracies exceeded the true accuracy in some folds, the minimum possible *p*-value was *0.002* (= 1/500). All tests were one-tailed.

The overall classification accuracy reported in the Results corresponds to the mean accuracy across the 28 folds for each participant. Reported box-plots in the figures indicate the lower and upper quartiles and maximum and minimum observed accuracy values (*n* = *28* folds per participant). Points shown beyond the whiskers of the box-plot do not adhere to the Tukey rule. No data points were excluded.

Biological replicates were the three individual participants (*n = 3*), each completing all sessions. Technical replicates were the repeated presentations of each sound within and across sessions (24 presentations per sound, across 8 sessions). Analyses were performed independently for each participant, and statistical tests were applied at the participant level unless otherwise noted.

## Data Availability

All source fMRI data and the associated timing files of sound presentations and button presses is publicly available via the *OpenNeuro* platform (openneuro.orgdatasets/ds006943/). Each participant’s structural image has been defaced to maintain their anonymity.

## Computer Code

All custom code used for analysis can be found in the publicly available *GitHub* repository https://github.com/DavidGHaydock/naturalSoundsPipeline

## Acknowledgements

This work was supported by the National Institutes of Health (R01DC004674), the National Science Foundation (SBE/BCS 2414066, 2420979), European Research Council (ERC) under the European Union’s Horizon 2020 research and innovation programme (grant agreement No. 740696), and in kind from the Birkbeck/UCL Centre for NeuroImaging (BUCNI).

## Materials & Correspondence

All materials used are publicly available via the links provided in the Data and Code Availability statements.

Correspondence to DH.

## Footnotes

1 Note that for clarity we use the labels that are provided with each of the parcellation schemes but are agnostic as to their correspondence with functional or anatomical regions or areas.

