## supplementary_figures for "Auditory Cortical Gradients Integrate Bottom-Up and Top-Down Structure During Natural Sound Categorisation"

### Supplemental Information

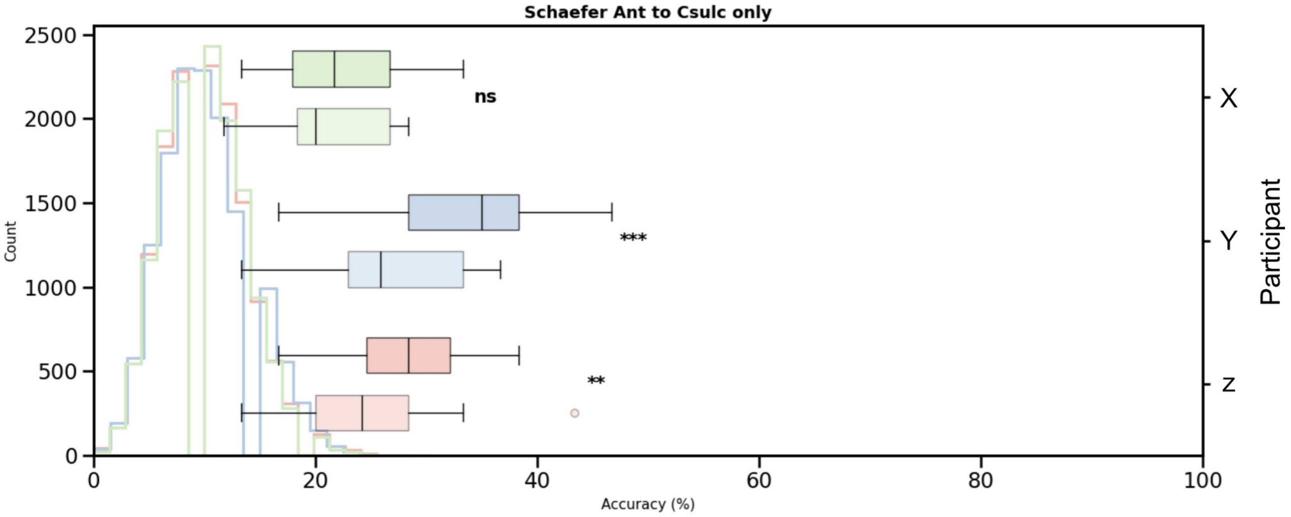


***Figure 1 - Classification Performance when only using frontal regions.*** *Classification performance of ROI- and gradient-based approaches using frontal region only. The frontal area of the "frontal-auditory subset" described in Figure 2 was used in classification, both with ROI- and gradient-based approaches. Box-plots show performance of classifier across folds, with histogram showing null distribution of accuracies across permutation test of 500 shuffles of sound category labels. Almost all observed fold accuracies fell within the null distribution of permutation test scores. T-test results between ROI and Gradient embedding approaches per parcellation per participant; * p<0.05, ** p<0.01, *** p<0.001 (ns = no significance).*

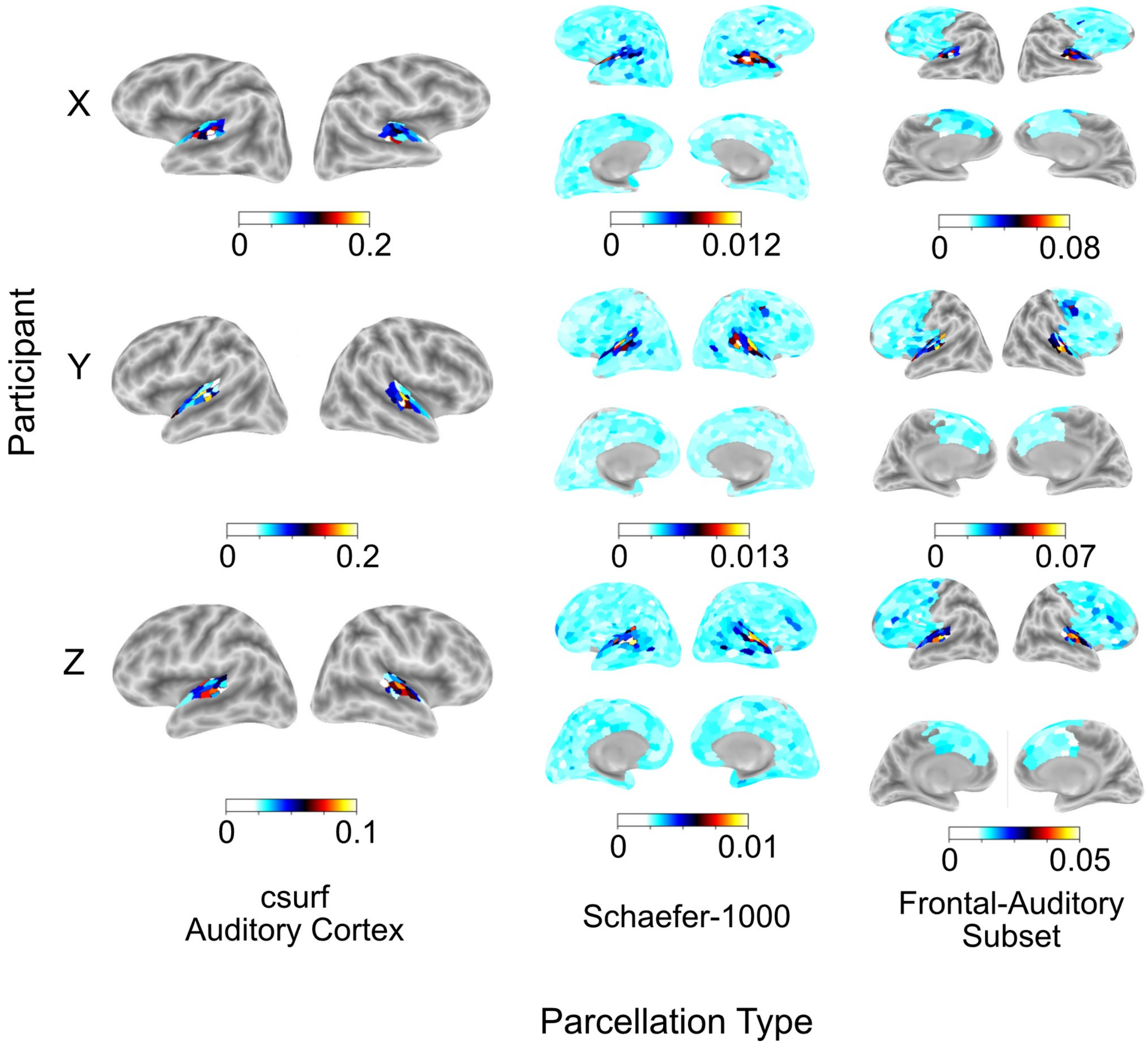


***Figure 2 - Feature importance maps of ROI data across participants for each parcellation type.*** *Each row shows a participant, and each column shows a parcellation type. Values on each ROI indicate feature importance. Values are percentages, with all values across parcels summing to 100%. Note the lack of importance values outside the auditory cortex in both the Schaefer-1000 and frontal-auditory subset parcellations.*

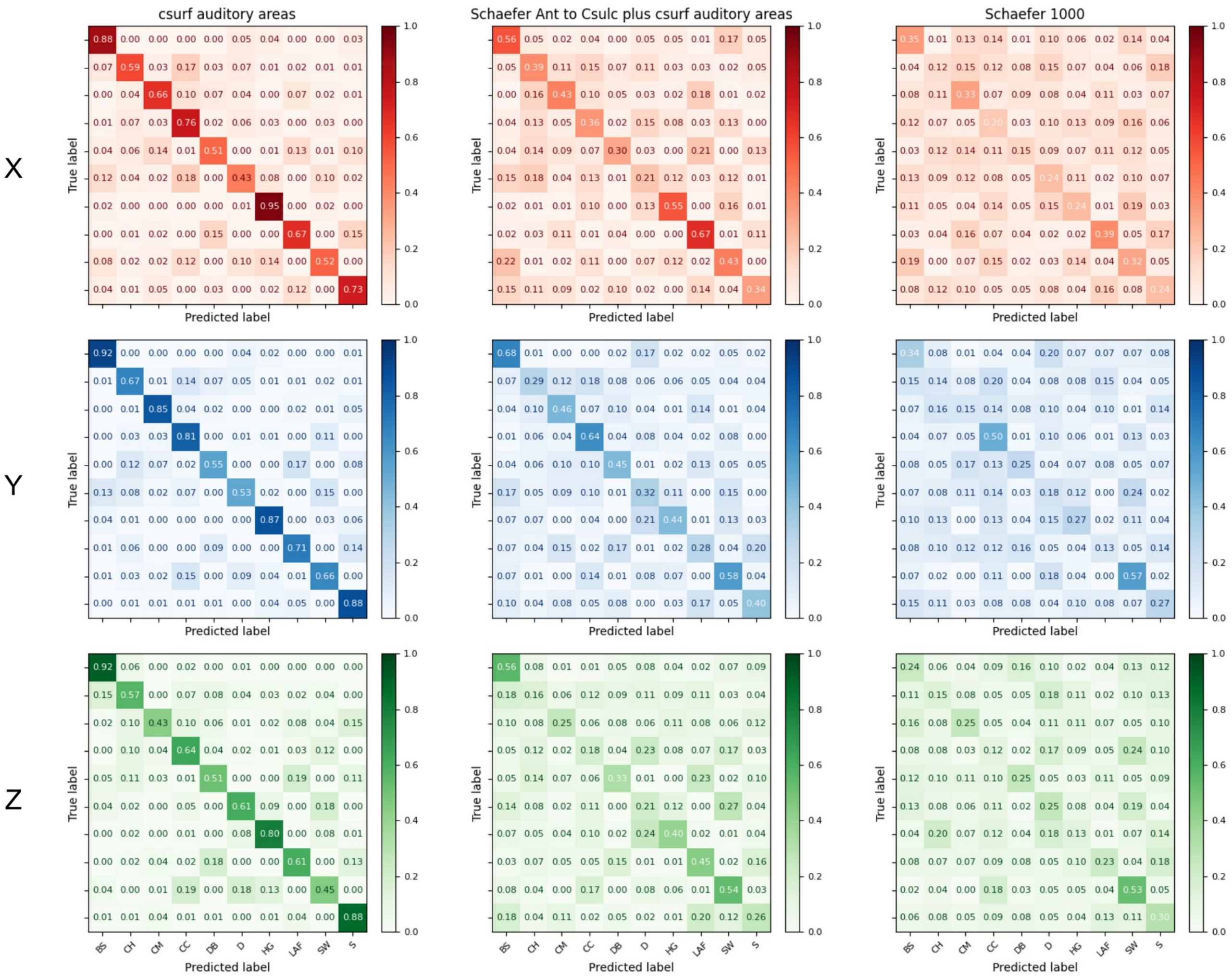


***Figure 3 - Confusion Matrices of all parcellations of gradient-space models across participants.*** *Each row corresponds to a participant, with each column corresponding to a parcellation type. First column is the equivalent to that shown in main Figure 3 in all participants. Second column shows same for frontal-auditory subset, and third for Schaefer-1000. BS – Bird Song; CH – Car Horn; CM – Cow Moo; CC – Crowd Cheering; DB – Dog Barking; D – Drilling; HG – Horse Galloping; LAF – Laughing Adult Female; SW – Sea Waves; S – Sneezing.*

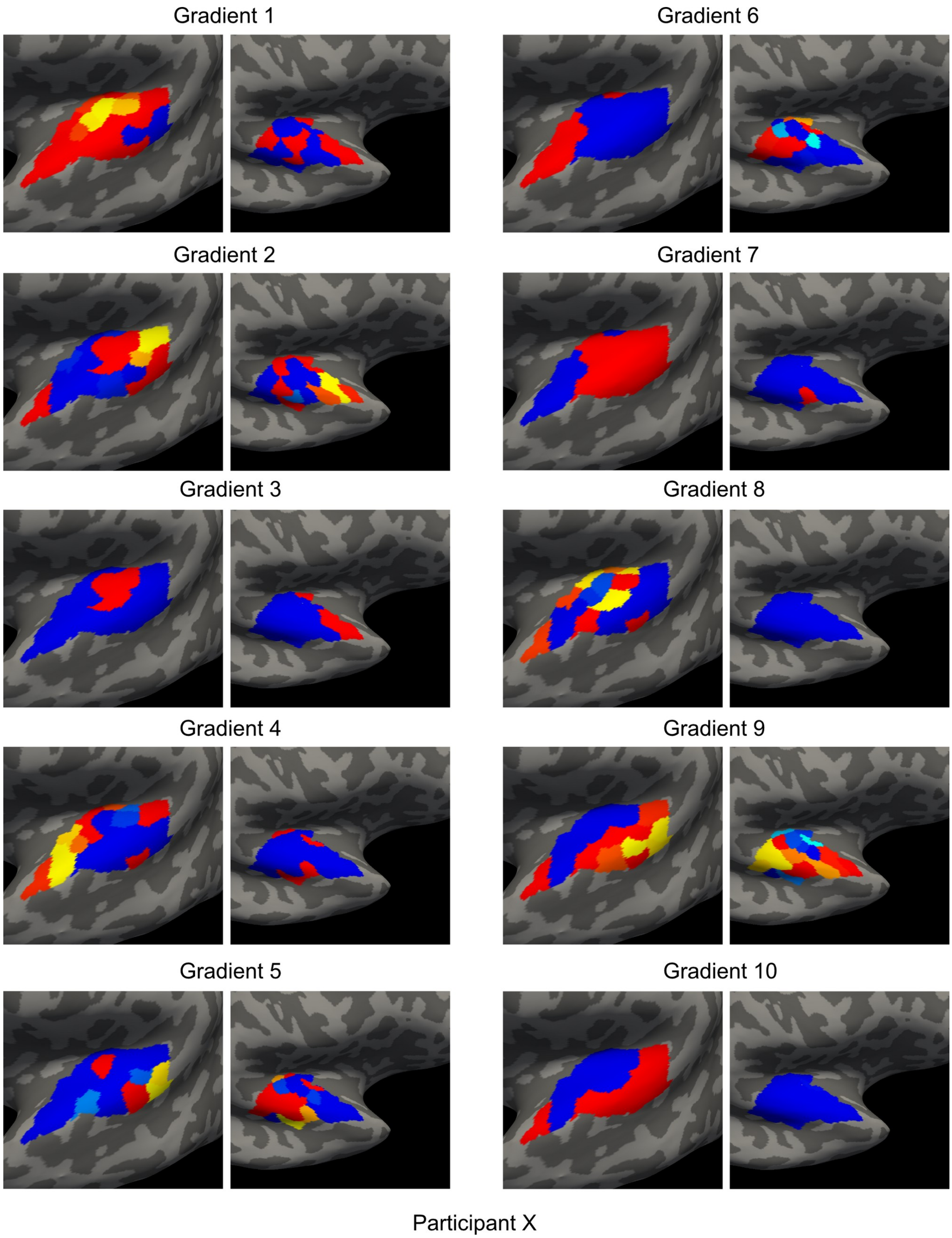


***Figure 4 - Participant X Auditory Cortex Gradient Maps on Individual Cortical Surface.*** *First 10 gradients for participant X in the auditory cortex parcellation. Given the number of parcels, images and .mgh files for all gradient maps can be found in the coda repository here:* https://github.com/DavidGHaydock/naturalSoundsPipeline

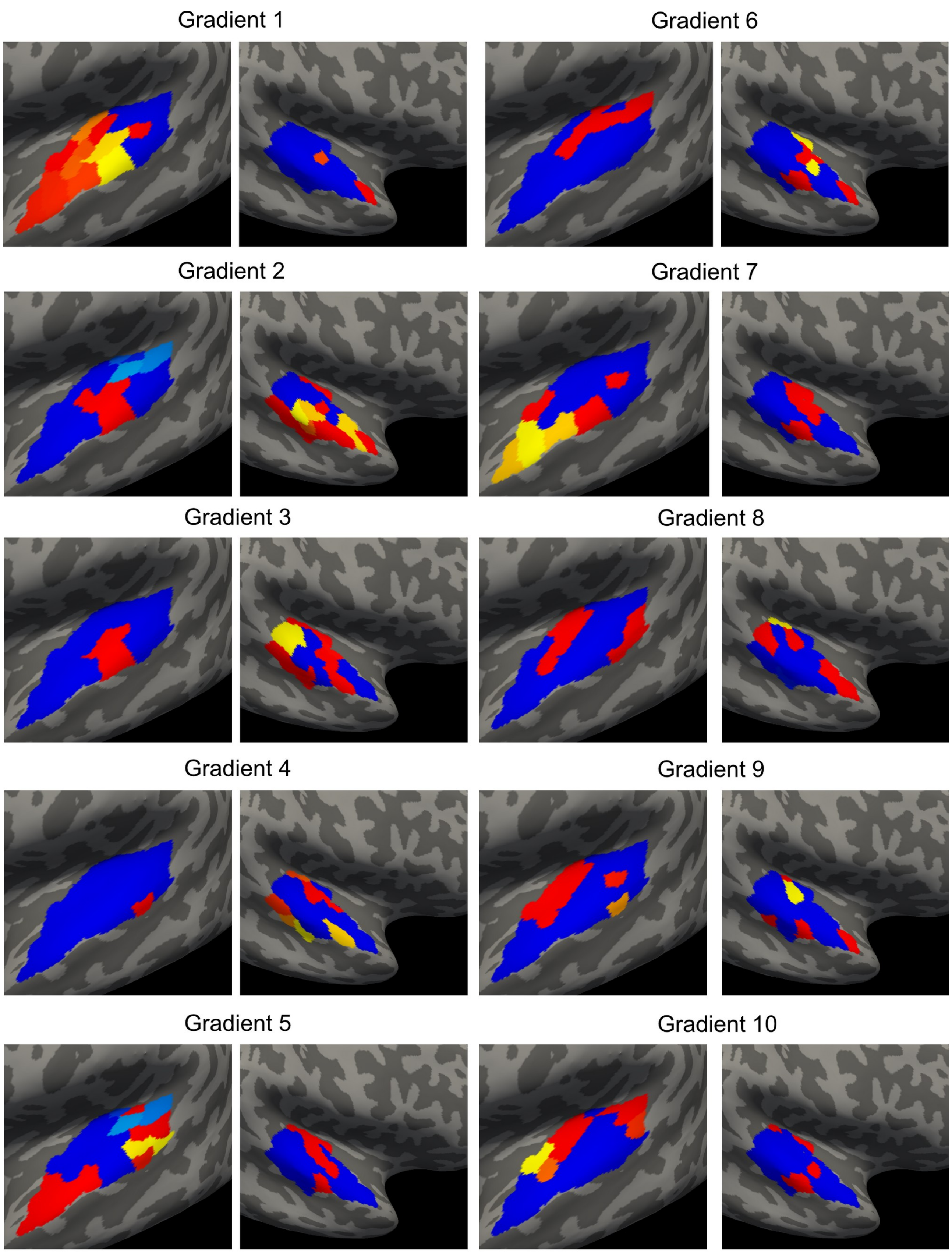


Participant Y

***Figure 5 - Participant Y Auditory Cortex Gradient Maps on Individual Cortical Surface.*** *First 10 gradients for participant Y in the auditory cortex parcellation. Given the number of parcels, images and .mgh files for all gradient maps can be found in the coda repository here:* https://github.com/DavidGHaydock/naturalSoundsPipeline

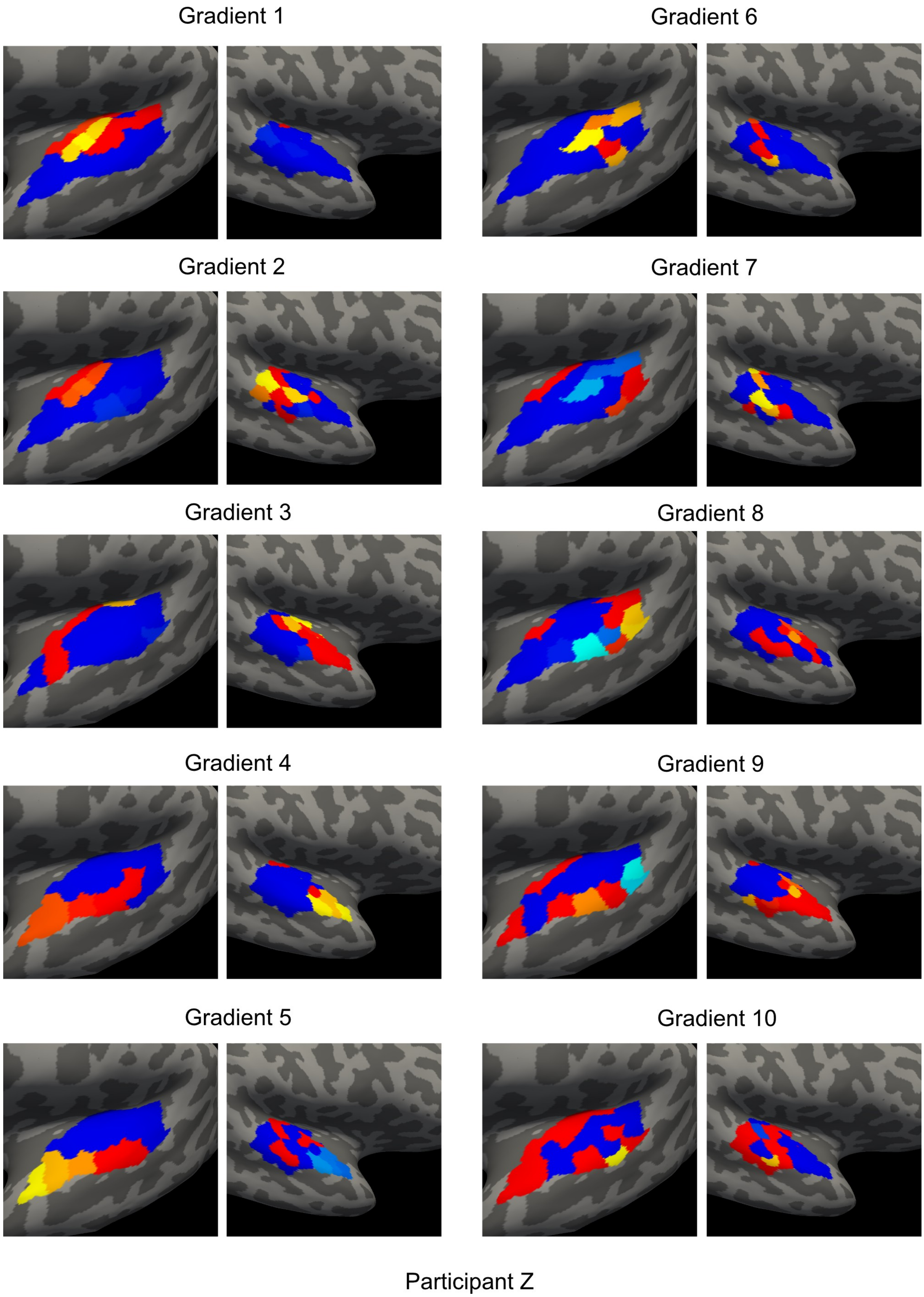


***Figure 6 - Participant Z Auditory Cortex Gradient Maps on Individual Cortical Surface.*** *First 10 gradients for participant Z in the auditory cortex parcellation. Given the number of parcels, images and .mgh files for all gradient maps can be found in the coda repository here:* https://github.com/DavidGHaydock/naturalSoundsPipeline

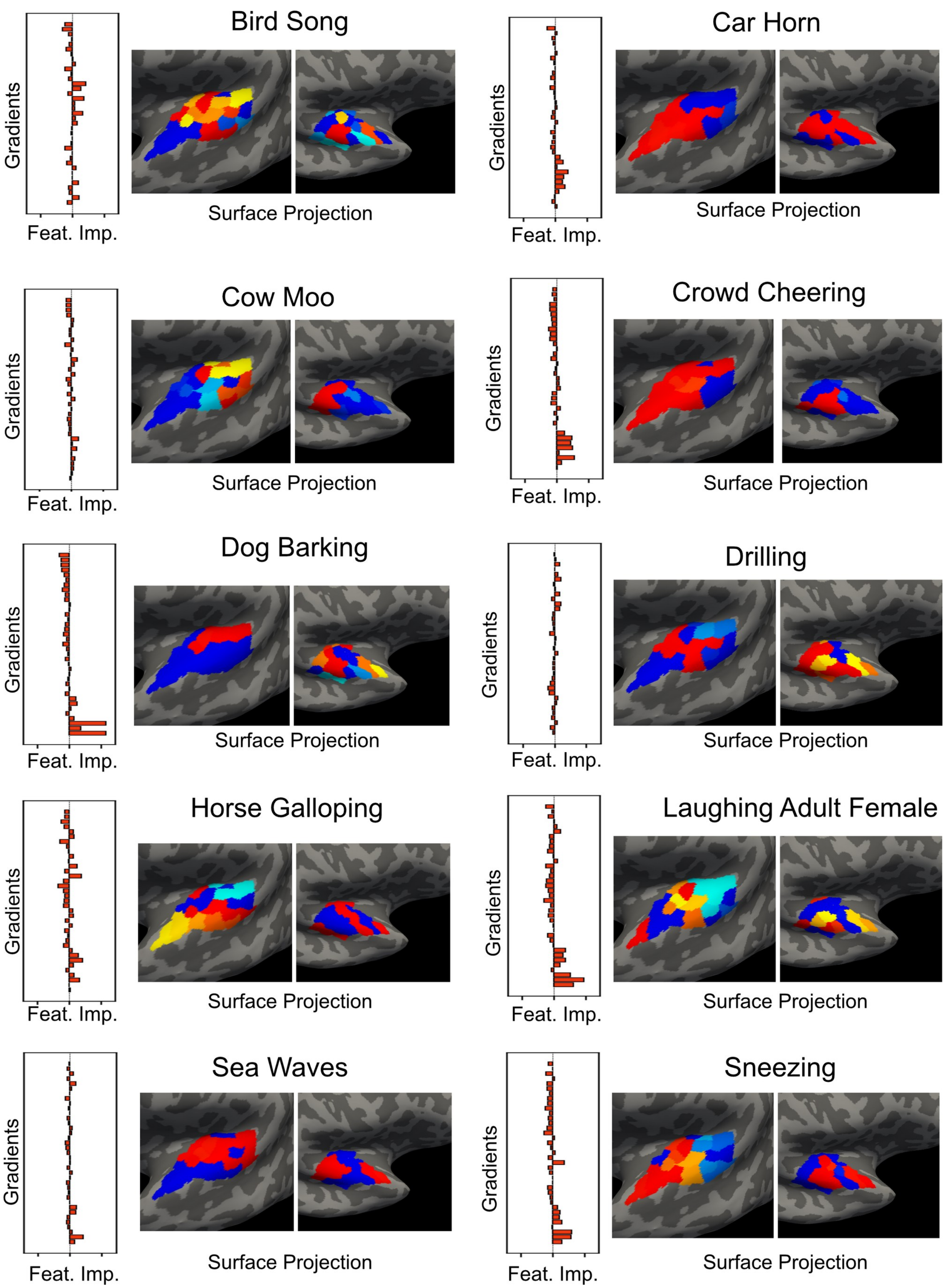


***Figure 7 - Participant X Gradient Space One-vs-Rest Category Feature Importance Projections.*** *Feature Importance of each OvR classifier per gradient, with surface projection of the given gradient weighting. Gradients are aligned over folds. (see Equation 1 for projection computation).*

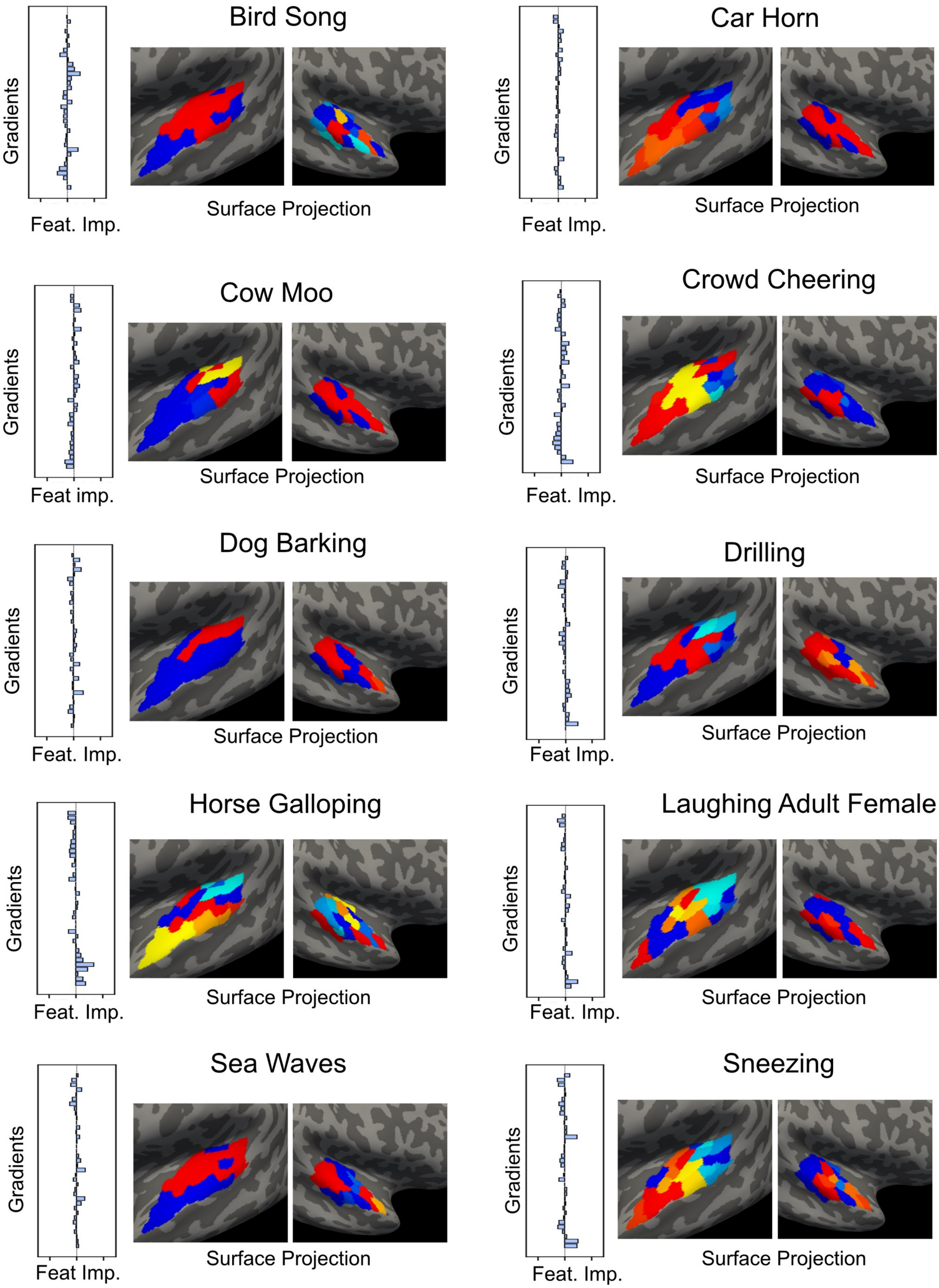


***Figure 8 - Participant Y Gradient Space One-vs-Rest Category Feature Importance Projections.*** *Feature Importance of each OvR classifier per gradient, with surface projection of the given gradient weighting. Gradients are aligned over folds. (see Equation 1 for projection computation).*

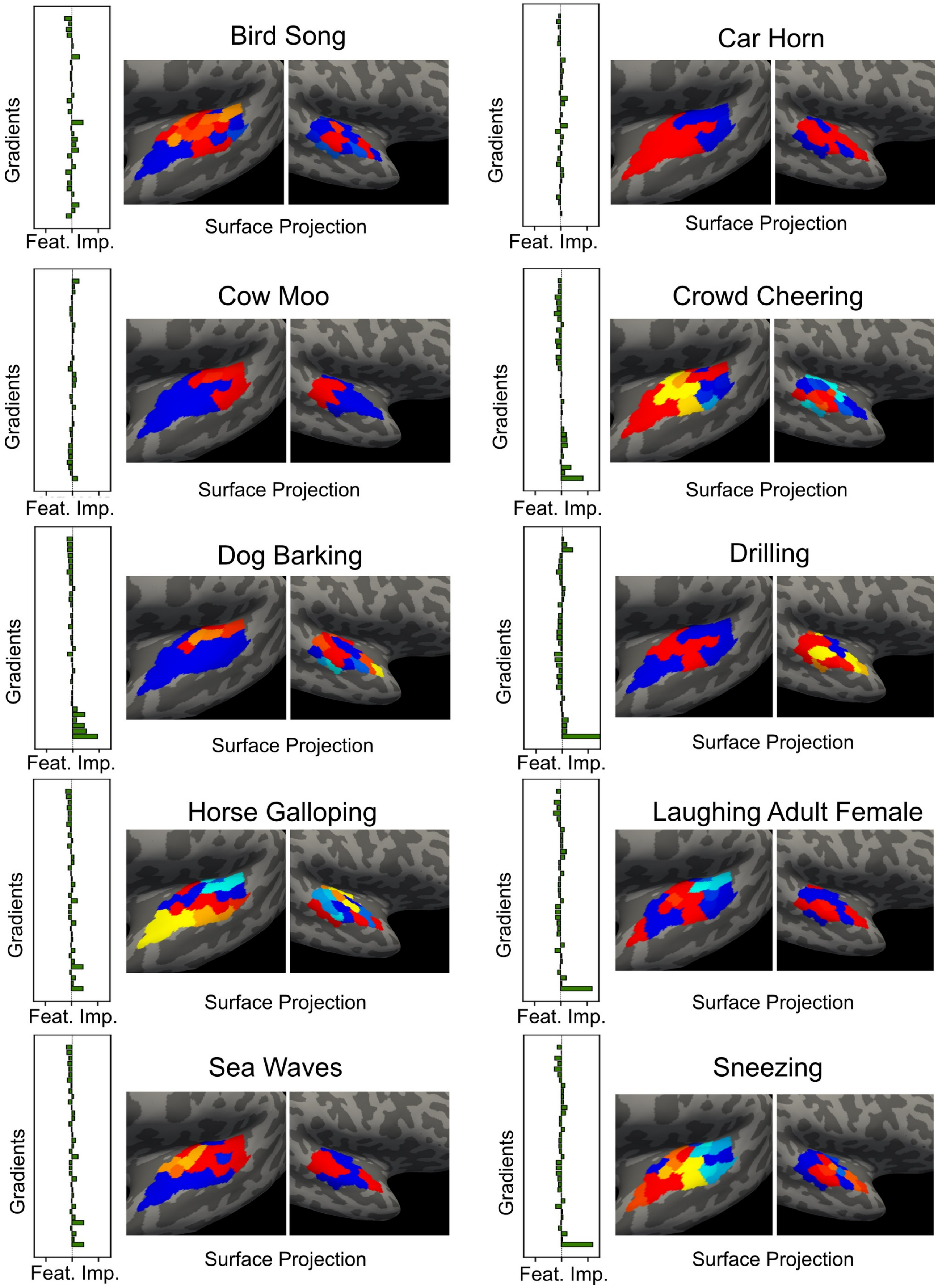


***Figure 9 - Participant Z Gradient Space One-vs-Rest Category Feature Importance Projections.*** *Feature Importance of each OvR classifier per gradient, with surface projection of the given gradient weighting. Gradients are aligned over folds. (see Equation 1 for projection computation).*

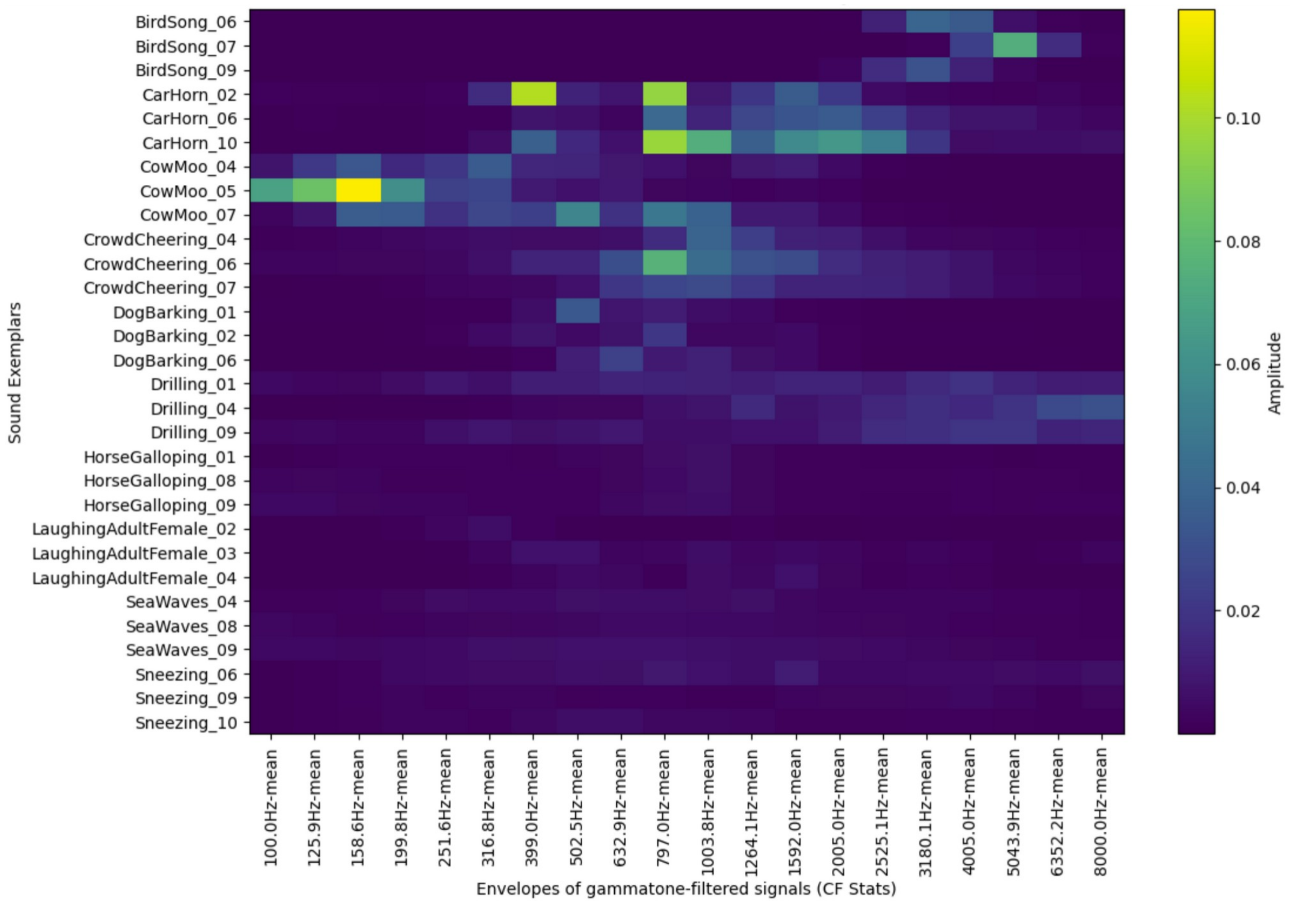


***Figure 10 - Gammatone mean representation of each sound exemplar.*** *Amplitude of mean for each filter used for input to PCA decomposition. The original names of each sound exemplar have been given here (note the numbers) so readers can cross-reference the sounds used with those in the dataset (Kachlicka, 2020), but the order of sound exemplars given here is equivalent to 1, 2, 3 in main text.*

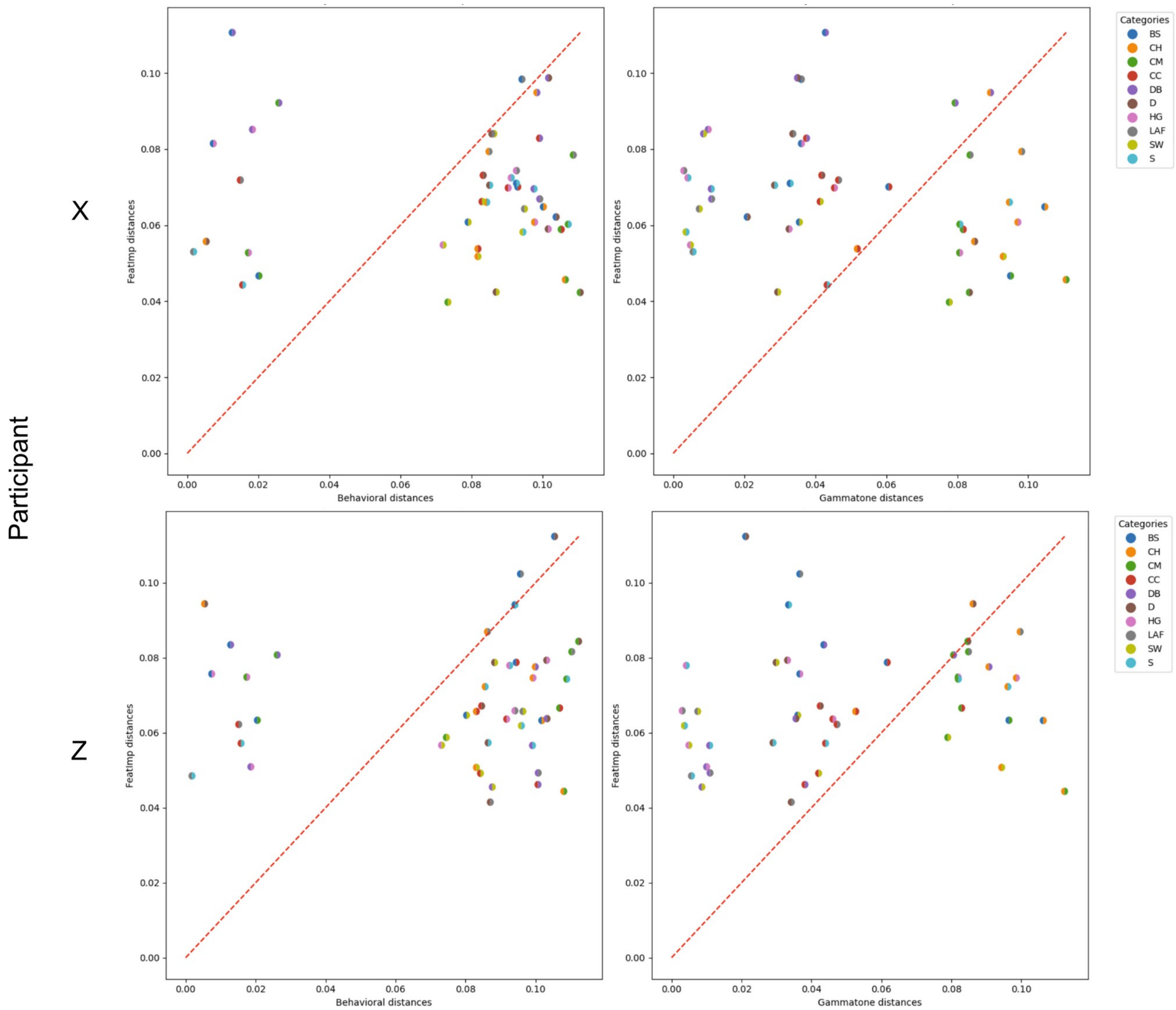


***Figure 11 - Participant Category RDM Comparison of gammatone and behavioural axes with gradient space for participants X and Z.*** *Difference in distance between sound categories for participant auditory cortex gradient model feature importance (y-axis) versus behavioural (left) and gammatone (right) distances for the same category pairs. Red mid-line shows where distances are equal. Participant Y equivalent is shown in main Figure 6. Note the similarity in category organisation across participants.*

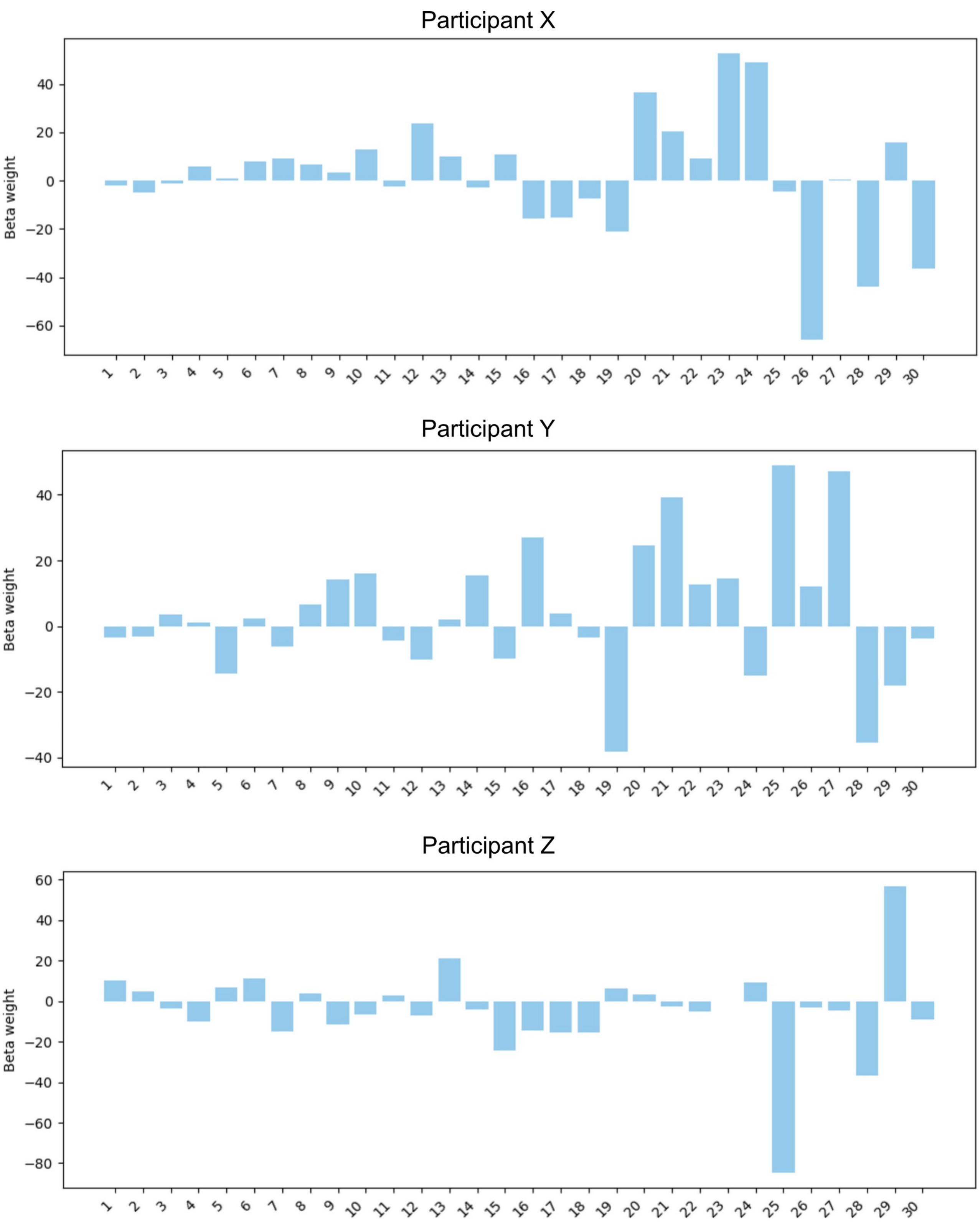


***Figure 12 - Participant tonotopy mapping onto individual gradient axes.*** *Weights for last 7 gradients are excluded due to abnormally large weights exploding the y-axis scale, making other weights hard to see. These large weights are likely due to some noise picked up in later gradients that explain less variance in the data than earlier ones.*
